# Trans-scale live imaging of an E5.5 mouse embryo using incubator-type dual-axes light-sheet microscopy

**DOI:** 10.1101/2023.05.01.538896

**Authors:** Go Shioi, Tomonobu M Watanabe, Junichi Kaneshiro, Yusuke Azuma, Shuichi Onami

**Affiliations:** Laboratory for Comprehensive Bioimaging, and; Laboratory for Developmental Dynamics, RIKEN Center for Biosystems Dynamics Research (BDR), 2-2-3, Minatojima-minamimachi, Chuo-ku, Kobe, 650-0047, Japan; Department of Stem Cell Biology, Research Institute for Radiation Biology and Medicine, Hiroshima University, 1-2-3 Kasumi, Minami-ku, Hiroshima, 734-8553, Japan

## Abstract

**We report our success in simultaneously tracking tissue formation and single-cell migration in a mouse embryo on embryonic day 5.5 for 12 h. The microscope, incubator, and observation protocols were comprehensively specialized and optimized to achieve this. The present system revealed new phenomena that had not previously been observed and promises to elucidate the mechanism of mouse embryonic development in the future.**

## MAIN

Because all body parts of a mouse, except for a section of the gut, consist of epiblast cell descendants,^1^ the developmental process of early mouse embryos must be meticulously spatiotemporally controlled. In particular, on embryonic day (E) 5.5, a cylinder-formed embryo exhibits the most primitive body axis, the anteroposterior (A-P) axis, which is the basis of subsequent morphogenesis.^2^ For A-P axis formation, asymmetric cell migration known as distal visceral endoderm (DVE) migration occurs over approximately 5 h,^3^ subsequently generating the chirality of embryo morphology and/or signaling cascade.^4, 5^ Long-term single-cell tracking in the epiblast during DVE migration with an observation of tissue formation is critical to unveil the mechanism by which the symmetry breaking of the cellular state causes symmetry breaking of the multicellular system during embryonic development. We succeeded in uninterrupted simultaneous tracking of single-cell migration and whole morphology change in a living E5.5-E6.0 mouse embryo, which has not previously been achieved.

The sample was an embryo of a mouse line that ubiquitously expressed green fluorescent protein-tagged Histone 2B (R26-H2B-EGFP mouse line).^6^ Because left–right asymmetry is formed after the E7.5 stage, we only needed to observe the hemisphere of an embryo to observe A-P axis formation. The required microscopic specifications were set as follows: a field of view of 0.3×0.3×0.15 mm^3^ for three-dimensional (3D) observation, a spatial resolution of less than 5 μm to individually resolve single nuclei, and a frame rate of 10 min to detect cell division for 12 h. To satisfy these specifications, we combined two branches of selective plane illumination microscopy (SPIM): multidirectional SPIM (mSPIM)^7^ to suppress shadowing artifacts, and dual-view inverted SPIM (diSPIM)^8^ for precise 3D rendering (Fig. 1a, b and Supplementary Note 1). Two multidirectional laser sheets were introduced to the embryo from both sides and rotated by 90° for dual-axes acquisition (Fig. 1a, *blue*, and Fig. 1c). Two image detectors were installed to acquire transverse/horizontal and longitudinal/vertical images sequentially (Fig. 1a, *green*).

**Fig. 1.**
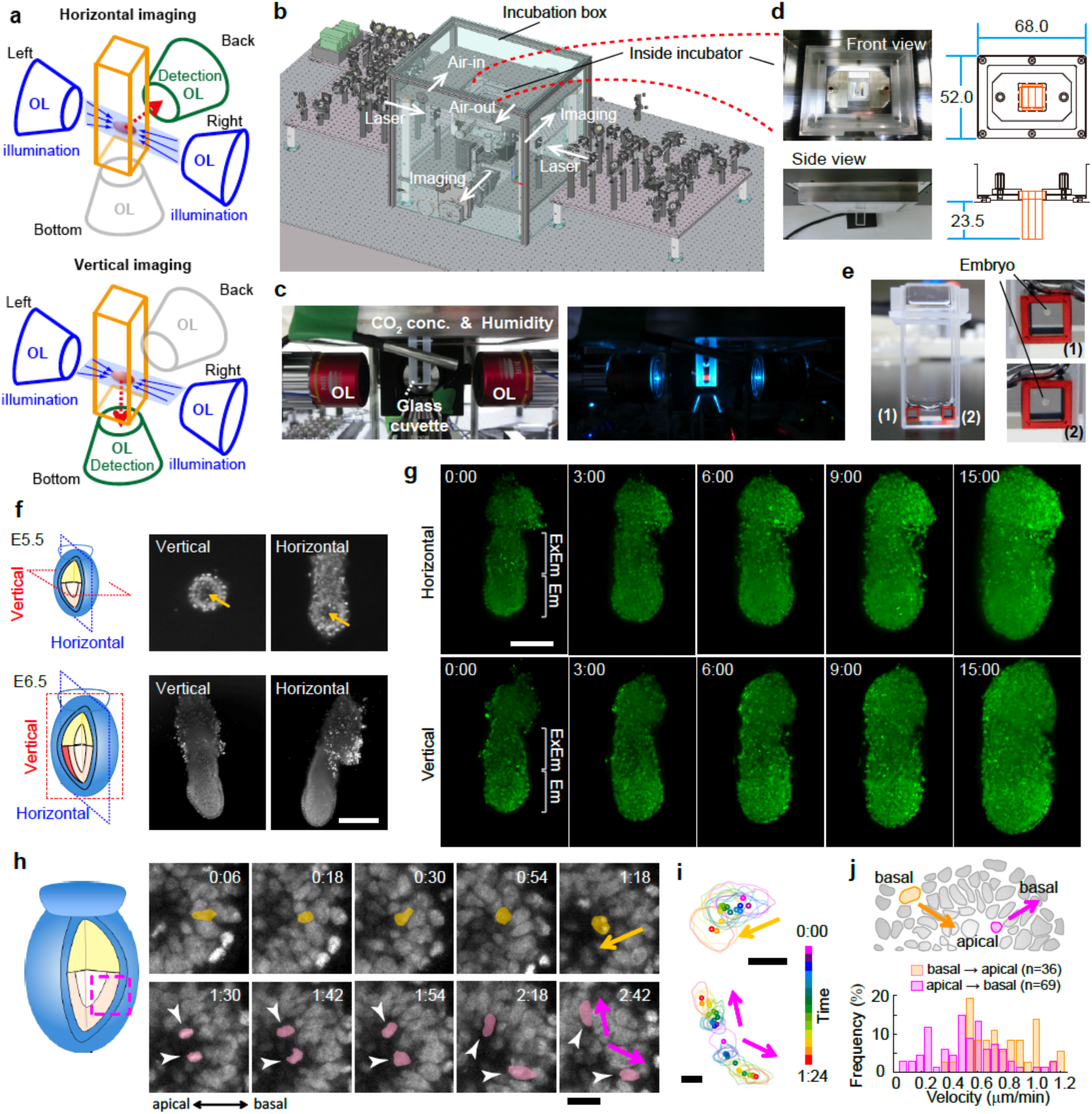
Microscopic system for in-toto single-cell observation of a mouse E5.5 embryo. (**a**) Schematic illustration of the placement of objectives for diSPIM. (**b**) Computer-aided design (CAD) illustration of the entire system. (**c**) Photograph of the inside of the incubation box. Left, under the light; right, during fluorescent imaging. OL, objective to produce a laser sheet. (**d**) Photograph and CAD illustration of the incubation chamber. The unit is mm. (**e**) Photograph of the glass cuvette (*left*) and collagen-filled cube (*right*). The cuvette, with a size of W12.5×L12.5×H30.0 mm for the outer dimension (W4.0×L11.1×H29.3 mm for the inner dimension), was composed of quartz. A 3 mm^3^ cube was filled with collagen-I gel and placed into the cuvette using a tweezer. (**f**) Typical example of dual-view imaging of an *R26-H2B-EGFP* mouse E5.5 (*upper*) and E6.5 (*lower*) embryo. Left, schematic; Middle, vertical image; Right, horizontal image. The scale bar is 100 μm. Arrows indicates the cavity. (**g**) Maximum intensity projection snapshots of the time-lapse images of an *R26-H2B-EGFP* mouse at the E5.5 stage. Forty-nine horizontal and 45 vertical images with a 5 μm z-step were acquired every 6 min for 15 h. The scale bar is 100 μm. Em, embryonic; ExEm, extraembryonic. (**h**) Snapshots of the time-lapse images of optical sections. A cell nucleus in the epiblast moves to divide near the proamniotic cavity (*yellow*), and each daughter cell intercalates into different positions in the epiblast (*magenta*). The scale bar is 20 μm. (**i**) Tracking result of cells in **h**. Lines, cell-outline; Dots, weight center. Upper, mother cell; lower, daughter cells. The scale bar is 10 μm. (**j**) Distribution of the velocity (µm/min) of INM-like movement from basal to apical (*yellow*, mean ± SD, 0.81 ± 0.24, n = 36) or from apical to basal (*magenta*, 0.56 ± 0.25, n = 69).

Only one group has previously achieved single-cell observation in a living E5.5 embryo; however, the duration of this observation was limited to 90 min,^9,10^ despite the successes of long-term observation before blastocyst^11, 12^ or after E6.5.^9, 13^ We speculated that the reason behind this was the instability of embryo culture on the microscope; while the culture conditions for the development of E5.5 embryos have been optimized in a floor-standing incubator,^14^ the temperature stability of the incubator is difficult to ensure on a microscope. Our incubation system had a two-layered structure on and covering the microscope (Fig. 1b, d), completely excluding the consideration of temperature instability (Supplementary Note 2). In addition, an embryo was embedded into a 3 mm cubic made of polycarbonate filled with collagen-I gel (Fig. 1e) to maintain normal developmental morphology of E5.5 embryos.^6^ The cube was fixed to the bottom of the cuvette via the surface tension of 150–200 μL of medium. Therefore, we successfully maintained normal development of a mouse embryo from E5.5 to E6.0, under fluorescent microscopic observation (Fig. 1f); a cell-less region called cavity were confirmed at the E5.5 stage (Fig. 1f, *yellow arrow*).

Another cause to limit observation duration is photodamage to the embryo due to continuous laser excitation. Importantly, reactive oxygen species (ROS) production, which is listed as the main mechanism of phototoxicity during fluorescent observation,^15^ did not correlate with the photodamage of cells in our case (Supplementary Note 3). We investigated photodamage by laser scanning using the growth rate of the embryoid body (EB) as the index and found that the interval time between continuous image acquisition did not necessarily reduce photodamage, and that maximizing the scan speed most effectively reduced photodamage (Supplementary Note 4). According to these findings, the exposure time for one image acquisition was set as short as possible (50 ms), and the three-directed light sheet of 0 ° and ± 6° was formed by scanning at maximum speed (16.7 ms for each directed sheet). The total number of frames for a 3D image was 300 frames along each axis. Moreover, we devised an optical setup to shorten the time for 90° rotation of the light sheets and adjusting the focus position (Supplementary Note 5). Finally, we observed all single cells in a hemispherical embryo every 6 min for 15 h (Fig. 1g, Supplementary Videos 1 and 2, and Supplementary Note 6) and confirmed the interkinetic nuclear movement (INM)-like migration previously reported^9^ (Fig. 1h, i). After an epiblast cell divided at the apical side close to the cavity (Fig. 1h, i, *yellow marks*), daughter cells intercalated into the different parts of the epiblast layer, and their nuclei moved toward the basal side (Fig. 1h, i, *magenta marks*). The mean velocity of INM-like migration in the epiblast was 0.81 ± 0.24 µm/min (n = 39) from basal to apical and 0.56 ± 0.25 µm/min (n = 69) from apical to basal (Fig. 1j), which are consistent with the values previously measured at the E6.5 stage,^9^ despite structural difference in tissue organization between E5.5 and E6.5. INM-like migration is thought to be driven by a fundamental mechanism regardless of the developmental stage.

In diSPIM, even when single-cell resolution was not achieved in either axis, it could be decomposed into the other axis, providing a definitive capture of cell division (Fig. 2a). This allowed us to perform statistic single-cell analysis. Daughter cells in INM-like migration were located apart after their nuclei reached the basal side (71 of 72 lineages, 98.6%) (Fig. 2b, c, *upper,* and Extended Data Fig. 1), whereas daughter cells in the visceral endoderm were localized close to each other (155 of 160 lineages, 96.9%) (Fig. 2c, *lower*). Considering the results of spatial transcriptomic heterogeneity in epiblasts along the distal-proximal axis at E5.25, E5.5, and E5.75,^16^ INM-like migration might contribute to relocating and shuffling cells to maintain cellular heterogeneity in a tissue or quickly expand the area of fate-determined cells. Further studies, combined with spatiotemporal transcriptomics, will reveal the true nature of this phenomenon.

**Fig. 2.**
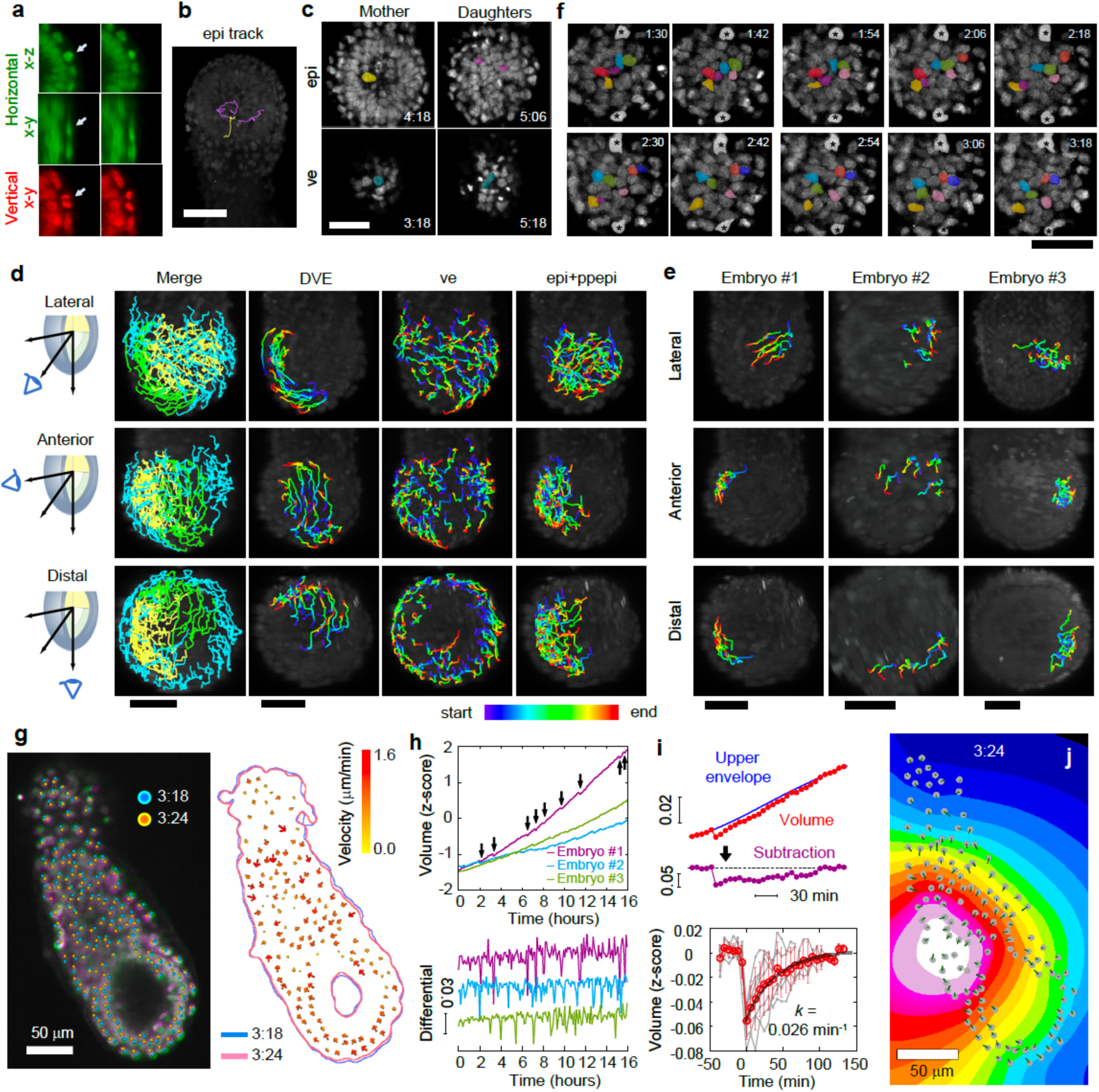
Trans-scale imaging of an E5.5 mouse embryo and analyses of cell movements. (**a**) Snapshots of horizontal (green) and vertical (red) images. The arrow indicates cell division realized in the vertical image, not in the horizontal image. (**b**) Traces of single lineage in the epiblast; a mother cell (*yellow*) and each daughter cell (*magenta*). (**c**) Front view of mitotic cells in the epiblast (*upper*) and visceral endoderm (*lower*). Left, mother cell; right, daughter cells. (**d**) Tracking results of individual cells during the DVE migration of embryo #1. Left panel (*merge*): green; DVE, blue; visceral endoderm, yellow; epiblast. (**e**) Tracking results of individual cells in the posterior-proximal region excluded from those of the epiblast of each embryo. Spectrum colors in **d** and **e** indicate elapsed time from the beginning (*blue*) to end (*red*) for 6 h. (**f**) Cell movement in the epiblast of an E5.5 mouse embryo. Colored, epiblast cell nuclei; asterisks, visceral endoderm cells. (**g**) Snapshot of shrinkage of an embryo. The left panel indicates that an embryo at time point 3:24 (*magenta*, elapsed time 204 min) is smaller than that at time point 3:18 (*green*, elapsed time 198 min). Cells at time points 3:18 and 3:24 are represented by cyan circles and orange circles, respectively. The right panel is the tracking results of individual cells. The color and size of an allow indicates migration velocity from 0.0 to 1.6 μm/min. The cyan and pink lines are outlines of the embryo at time points 3:18 and 3:24, respectively. (**h**) Time course of the volume of an embryo during live imaging. The arrows show the timing of shrinkage. Upper, the z-score, the standard deviation from the mean; lower, differential of the upper panel. The colors indicate each embryo, that is, #1 (*magenta*), #2 (*cyan*), and #3 (*light green*). (**i**) Hiccup-like behavior after eliminating entire growth. A time course of hiccup-like behavior was obtained by subtraction of the upper envelope from volume data (*upper*). The seven time-courses could be extracted (*gray lines*). The traces were set from the moment of shrinkage to time zero and averaged (*red*). Black line, fitting results by single exponential decay, *f*(*t*) = *a*·exp (-*k*·*t*); error bars, standard deviation. (**j**) Visualization of a shrinkage center of embryo #1 at time point 3:24 from the horizontal view. Colors indicates a shrinkage level at 4-bit scale, from white (min) to black (max). The gray circles are cell positions of the embryo and green lines indicates distances of cell migration within a frame. The scale bars are 50 μm.

Because all single cells in a hemispherical embryo are isolated, the cells could be classified into the epiblast, visceral endoderm, and DVE according to their positions and movement (Fig. 2d). The collective migration of the DVE was successfully identified in all three embryos (Extended Data Fig. 2, *DVE*). During DVE migration, several epiblast cells collectively moved in an embryo (Fig. 2e, *#1*) and moved from the prospective posterior-proximal side to the prospective anterior-distal side close to the visceral endoderm (Fig. 2f, *colored*), while the visceral endoderm cells hardly moved (Fig. 2f, *asterisks*). Because the collectivity of epiblast migration in embryo #1 was eliminated with the cessation of DVE migration, the epiblast cells were thought to be passively moved by DVE migration. Meanwhile, we did not observe collective cell movement in the two other embryos (Fig. 2e, *#2 and #3*). In addition, collective movement of visceral endoderm cells in the lateral region, as previously reported,^6^ was observed in embryos #1 and #2, but not clearly in embryo #3 (Fig. 2d and Extended Data Fig. 2, *ve*). In short, epiblast and visceral endoderm cells seemed to passively move to resolve the distortion of cell arrangement caused by active DVE migration. A large-scale analysis must be performed in the future to determine the authenticity, biological meaning, and mechanism of these variations among individual embryos using the present method.

Additionally, we found an interesting behavior in the embryo volume: the embryos grew monotonously but often shrunk abruptly, resembling a hiccup (Fig. 2g). Hiccup-like behavior was observed in all three embryos analyzed (Fig. 2h and Extended Data Fig. 3), and its occurrence frequency was 0.019 ± 0.017 min^-1^ (n = 27). The volume recovered exponentially after shrinkage, and its recovery rate was estimated to be 0.026 min^-1^ by an exponential approximation (Fig. 2i). The spatial- and time-scales of the hiccup-like behavior were similar to the abrupt shrinkage of a cavity until the hatching of a pre-implantation embryo, that is, a blastocyst.^17^ The cavity in an embryo is thought to work on tissue self-organization as not only a mechanical cue but also a biochemical signaling source, while the various signal molecules work on cell proliferation and differentiation at the E5.5 stage.^4, 18^ However, because the cavity was not the shrinkage center at E5.5 (Fig. 2j), the hiccup-like behavior we observed was distinguished from the abrupt shrinkage in a blastocyst. Although the same embryo exhibited the same shrinkage center even at different time points (Extended Data Fig. 4a, b), the shrinkage center depended on the embryo, but was limited to the periphery or outside of the amniotic cavity. (Extended Data Fig. 4c, d). Osmotic pressure in the cavity affects embryo size and cell fate,^19^ and external mechanical stress to an embryo promotes its A-P axis formation.^20^ It is possible that abrupt shrinkage causes an osmotic difference between the cavity and the outside of the embryo to regulate mechanical signaling in a post-implantation embryo. Hence, we hypothesize that the hiccup-like behavior in an E5.5 embryo may generate chirality of the mechanochemical signals, leading to position-dependent cell fate for A-P axis formation. Although an investigation into the relevance of our hypothesis remains a future issue, this study shows that the present microscope system visualizes each cell migrating toward specific sites, that is, one side of the extraembryonic region, for the first time, providing new insight into A-P axis formation.

In conclusion, we achieved simultaneous long-term tracking of single cells and tissue formation dynamics in a hemispherical embryo for over 12 h at the E5.5 stage. More data on embryos are required to verify the phenomena observed in this study and to understand the mutual/causal relationships between them. There is no doubt that single-cell analysis technology using deep learning will develop further; however, applying deep learning requires a large amount of data. The present microscope system is now available for use. We hope that more people will use this system in combination with deep learning to fully understand multi-layered symmetry breaking in A-P axis formation.

## ACKNOWLEDGMENTS

We thank Mari Kaneko and Hiroshi Kiyonari (RIKEN) for preparation of the ES cells and LARGE for animal housing and resources. This study was supported by MEXT Grant-in-Aid for Scientific Research on Innovative Areas “Singularity Biology,” Grant Number JP18H05409. We would like to thank Editage (www.editage.com) for English language editing.

## AUTHOR CONTRIBUTIONS

G. S. performed all experiments and co-wrote the manuscript. T. M. W. supervised microscope construction and mainly wrote the manuscript. J. K. constructed the microscope system and the software for device control. Y. A. contributed to the image registration and embryo volume calculations. S. O. led and designed the study.

## DATA AVAILABILITY

All videos used in the main results are available on the following site: https://doi.org/10.24631/ssbd.repos.2022.07.243. The other videos and analyses results can be made available from the corresponding author upon reasonable request.

## Online Methods

### Animals

*R26-H2B-EGFP* mice (accession no. CDB0238K: http://www2.clst.riken.jp/arg/reporter_mice.html)^21^ were used. The animals were housed in environmentally controlled rooms, and all experimental procedures involving animals were reviewed and approved by the Institutional Animal Care and Use Committee of the RIKEN Kobe Branch.

### Embryo culture

Embryos were harvested and cultured as previously described.^22, 23^ In brief, embryonic day 5.5 (E5.5) or E6.5 embryos were cultured in collagen gels (Nitta Gelatin Cellmatrix TypeI-A), which were mixed with the embryo medium (Dulbecco’s Modified Eagle Medium (DMEM) (Sigma Aldrich D2902) supplemented with 50% rat serum, 1 mM of β-mercaptoethanol (Sigma Aldrich M-3148), 1 mM of sodium pyruvate (Gibco 11360-070), and 100 μM of nonessential amino acids (Gibco 11140-050)). The embryos were cultured in a controlled environment at 37 °C and 5% CO_2_. A cultured embryo under suitable irradiation conditions for image acquisition exhibited as good a growth as a similarly mounted embryo on the opposite side of the cuvette.

### Culture environment

A chamber and an incubation box were developed in this study. The temperature in the incubation box, including the chamber, was controlled at 37 °C using a heater (Tokken TK-0003HU20). The concentration of CO_2_ was controlled at 5% using a gas mixture (Tokken TK-MIGM01-02) only in the chamber. To achieve high humidity, 5% CO_2_ gas was passed through in water heated at 37.5 °C with a bubbling system (Tokken TK-HE05). The detail is described in Supplementary Note 2.

### Embryoid body formation

TT2 embryonic stem (ES) cells^24^ on feeder cells cultured in ES medium (DMEM (Gibco 12100-046) supplemented with 1000 units/ml of leukemia inhibitory factor (LIF) (MERCK ESG1107), 1 mM of β-mercaptoethanol (Sigma Aldrich M-3148), 1 mM of sodium pyruvate (Gibco 11360-070), 100 μM of Modified Eagle Medium (MEM) non-essential amino acids (Gibco 11140-050)), and 10% fetal bovine serum (FBS) until the ES cells were 70–80% confluent. The ES cells were collected and suspended in ES medium without LIF (LIF-ES medium) and seeded into a low-adhesion U-shaped-bottom 96 well plate (SUMITOMO BAKELITE MS-9096U) at a density of 500 cells per well for photodamage, 1000 cells per well for reactive oxygen species (ROS) production experiments, and 2000 cells per well for temperature experiments. Embryoid bodies formed after two days were used for the photodamage experiment and after a day for the ROS production and temperature experiments.

### Quantification of photodamage

Embryoid bodies in LIF-ES medium were mounted on a glass-bottom dish (Matsunami D11040) and set on the stage of the microscope. The samples were irradiated with laser illumination in single scanning mode. Irradiated embryoid bodies were returned to the wells and cultured independently. A Nikon TiE-A1RSi confocal laser-scanning microscope with a 20-fold objective lens (Nikon CFI Plan Apo VC 20×) was used.

### Growth ratio of embryoid body

The cross-sectional area of the image of an embryoid body was measured before (Area.before) and after the embryoid body was irradiated. The area was measured again 24 h later (Area.after). The ratio of Area.after to Area.before was calculated as the embryoid body growth ratio. Images of the embryoid bodies were acquired using a Nikon ECLIPSE TS100 equipped with a 4-fold objective lens (Nikon CFI Plan Apo 4×) and a CCD camera (Nikon DS-Fi1).

### Image acquisition by viSPIM

A mouse embryo was embedded in a cube (NK system CIDH-29) containing collagen gels mixed with embryo medium. The cube was then placed at the bottom of a customized glass cuvette with 150–200 μL of embryo medium. The cuvette was set on the chamber, and images were acquired every 5 or 6 min with a 10-fold objective lens (Olympus UPlanFL N 10×) or 20-fold objective lens (OptoSigma Plan Apo L 20×), 488 nm laser (Kyocera, JUNO488), band-pass filter (Semrock FF01-525/45-25), and complementary metal oxide semiconductor (CMOS) camera (Hamamatsu Photonics ORCA-Flash4.0).

### Image registration

To facilitate the visual recognition of each nucleus, we fused the images acquired in the horizontal and vertical directions. First, the image resolutions in the xy- and z-directions were equalized. Because the resolutions were 0.65 and 5.0 μm/pixel in the xy- and z-directions, respectively, the number of z-slices were increased by 5.0/0.65 times via bicubic interpolation. Equalization was applied to the horizontal and vertical image stacks at each time point. Second, the image orientations were aligned by rotating either of the stacks in three dimensions. Third, the image stack was registered by spatial shifting, and thus “translation.” The image registration techniques are explained elsewhere,^25^ although we used Mattes mutual information^26, 27^ as the metric and the (1+1)-evolution strategy^28^ as the optimizer to compute the transformation function. The function was computed for the image stack at the initial time point and applied to those at all time points. Finally, the registered image stack was added to the image stack in the opposite direction at each time point to create a fused image stack. If the horizontal image stack was registered, it was added to the vertical image stack and vice versa. Therefore, two fused image stacks were created at each time point. The resulting image stacks were used for manual tracking.

### Volume computation

To compute embryonic volume, the embryonic region was segmented at each time point. First, the image stack was denoised by applying a three-dimensional (3D) Gaussian filter (*σ*=0.5 [pixels]). The denoised image stack was subsequently binarized via global thresholding, where the threshold was determined by multiplying the mean intensity of the stack by 1.2. If holes were present in the foreground, they were filled. Embryonic volume was obtained by measuring the number of foreground pixels and converting them to the actual scale. The shrinkage timings were determined as outliers of the volume ratio between two adjacent time points based on three-sigma rule.

### Visualization of the location of shrinkage

We visualized each coordinate of the images as a point where the cells came close to or apart from the abrupt shrinkage. A vector from a certain position *P* to a certain cell *C_i_*, where *i* is the cell number, at a certain frame *t* is denoted *PC_i_* (*x_i_*_,*t*_, *y_i_*_,*t*_). A pixel value of a new image at a frame *t*, *I*(*x_t_*, *y_t_*), was the summation of inner products between *PC_i_* (*x_i_*_,*t*_, *y_i_*_,*t*_) and *PC_i_* (*x_i_*_,*t*+1_, *y_i_*_,*t*+1_) normalized their scalar, as followed,

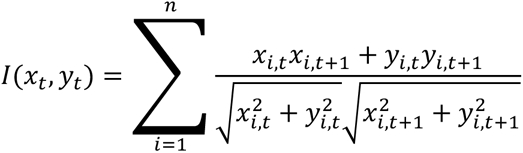

, where *n* is the number of the observed cells. For visualization, *I*(*x*, *y*) values were plotted on an inverted 8-bit grayscale, and visualized using a 16-colors look-up-table.

### Image analysis

Maximum intensity projection images of time-lapse images were obtained using MetaMorph software (Molecular Devices). Cell tracking analysis was performed manually using Imaris software (Oxford Instruments).

## Extended Data Figures

**Extended Data Fig. 1.**
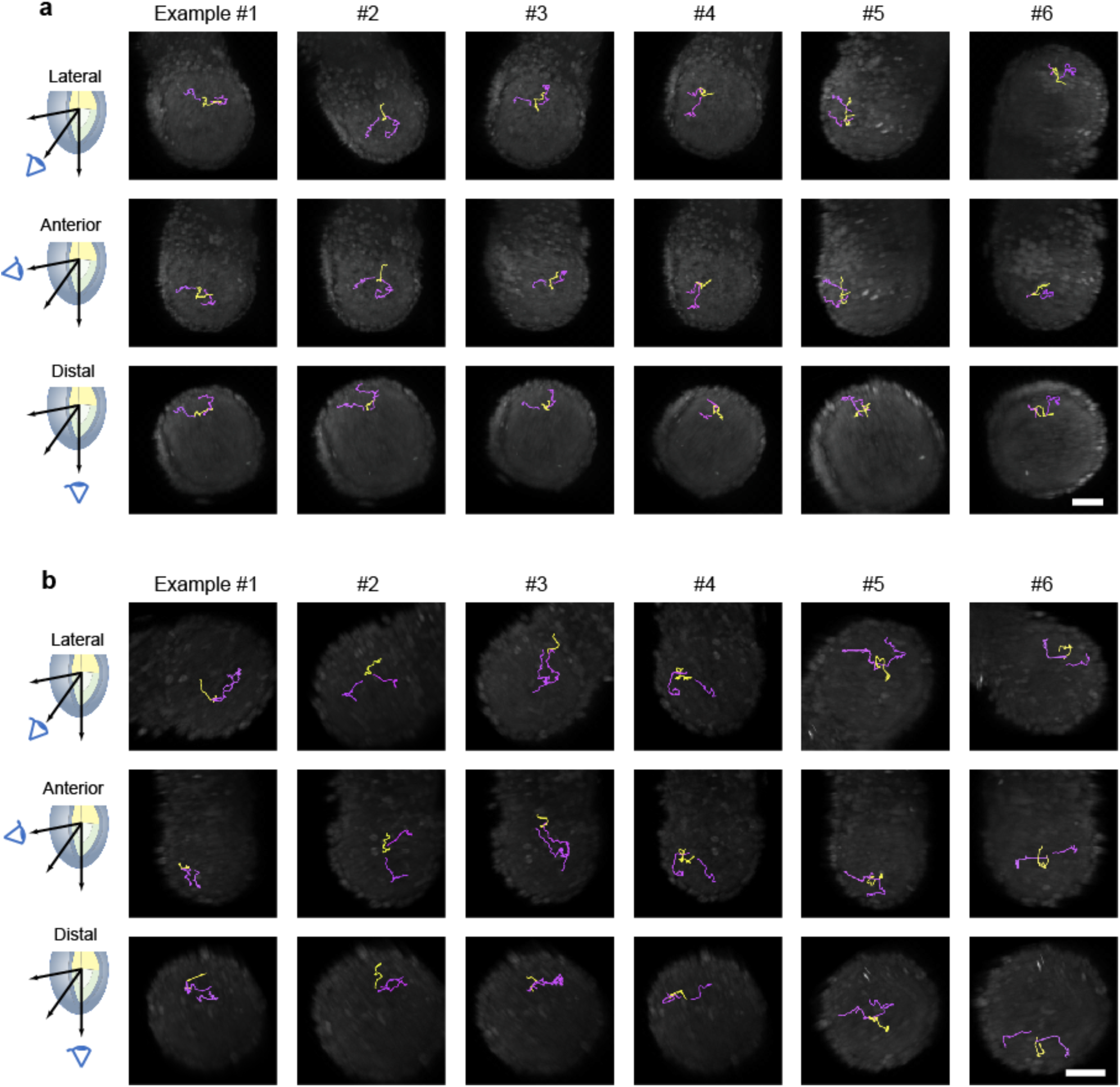
Examples of the migration of a single lineage in the epiblast. (**a, b**) Typical six traces of the cell movement of a mother (*yellow*) and each divided daughter cell (*magenta*) in embryo #1 (**a**) and #3 (**b**). The scale bars are 50 μm.

**Extended Data Fig. 2.**
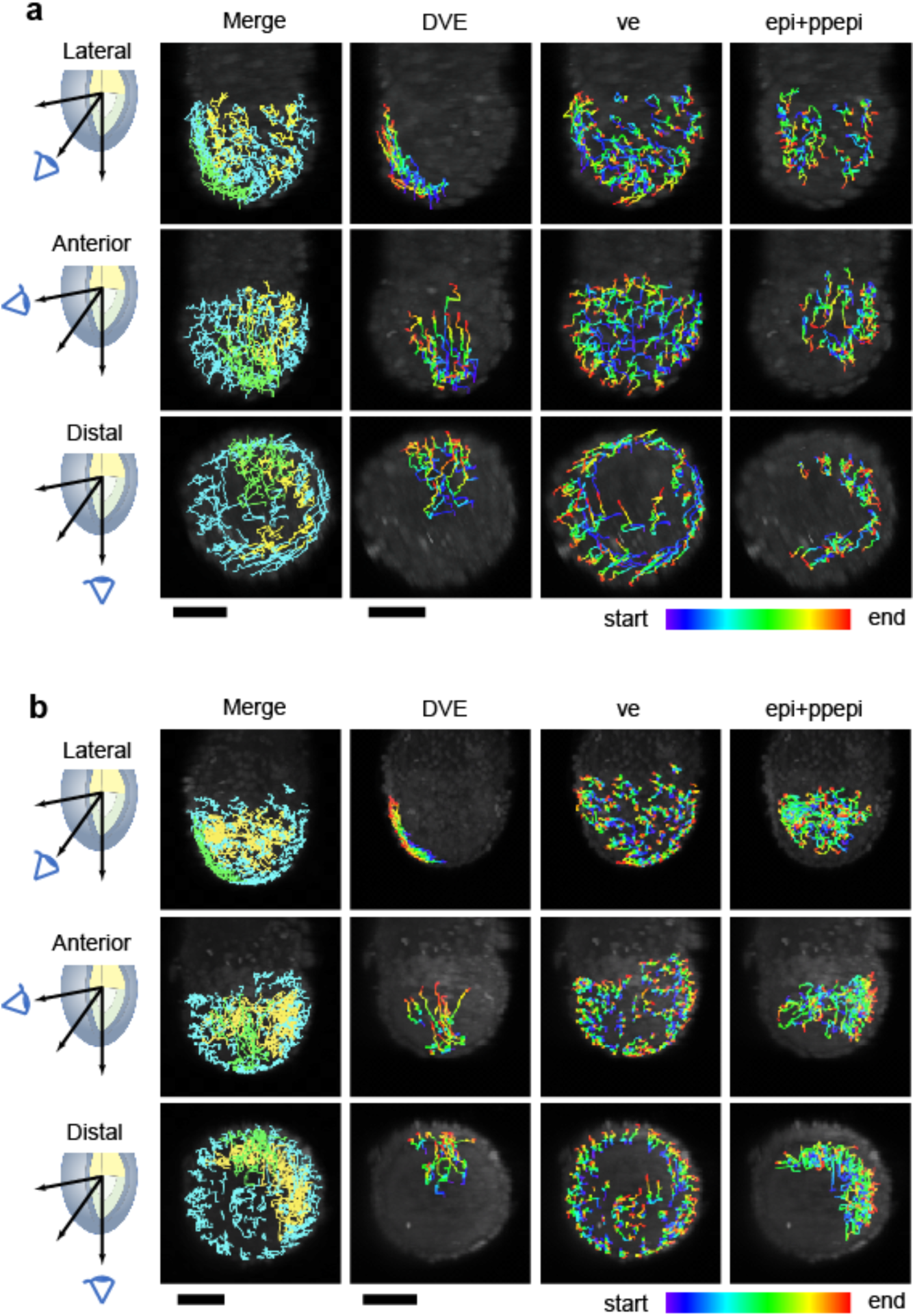
In-toto single-cell tracking in an E5.5 mouse embryo. Tracking results of single cells during distal visceral endoderm (DVE) migration of embryos #2 (**a**) and #3 (**b**). Left panel: green, DVE; blue, visceral endoderm; yellow, epiblast. Right three columns: cell tracks in each region shown by the spectrum color from start (*blue*) to finish (*red*). The scale bars are 50 μm.

**Extended Data Fig. 3.**
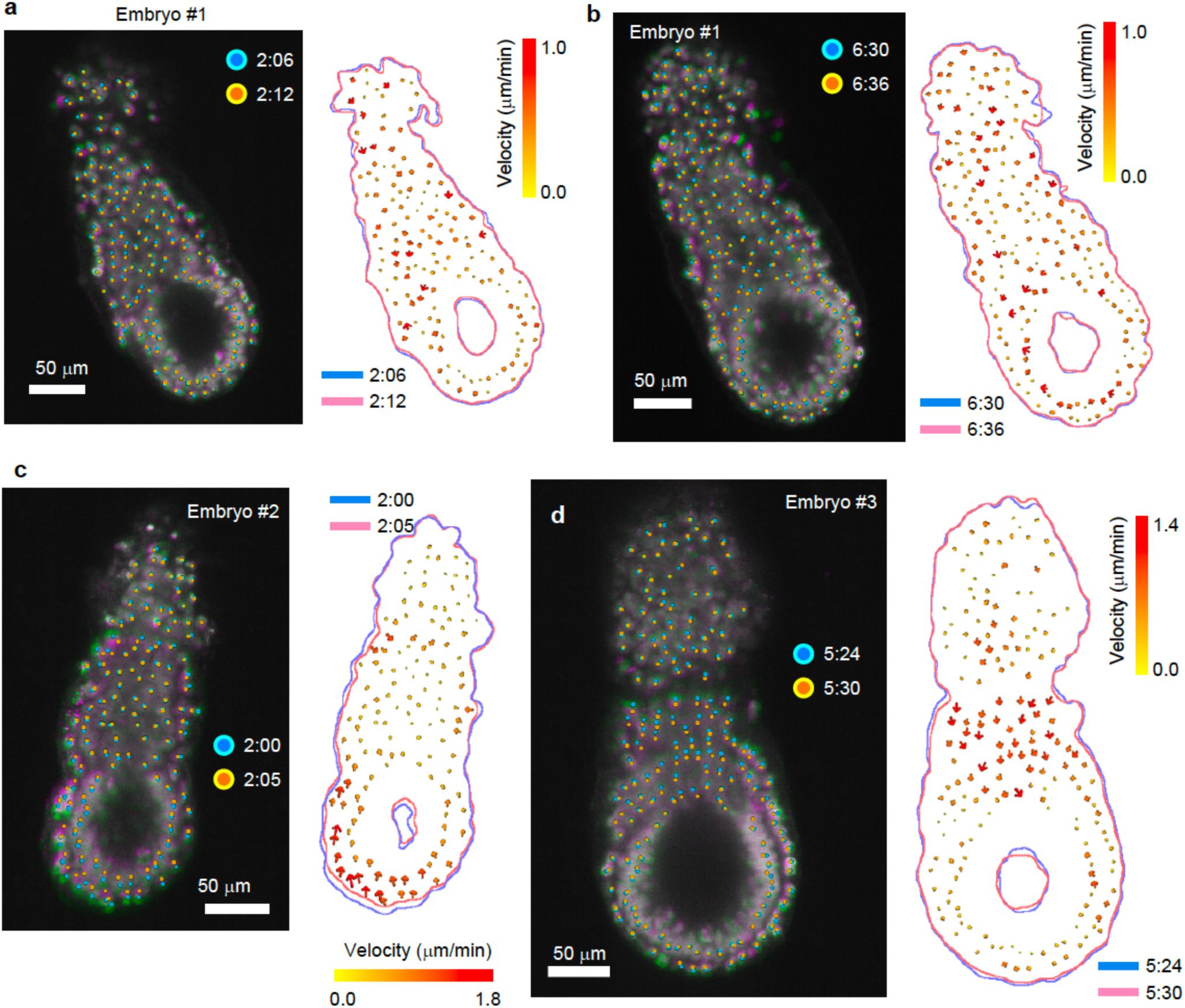
Typical example of hiccup-like behavior in an E5.5 mouse embryo. (**a–d**) Snapshots of the shrinkage of embryos #1 (**a, b**), #2 (**c**), and #3 (**d**) at two consecutive time points (*first, green; second, magenta*). Cells at the first and second time points are shown as cyan circles and orange circles, respectively. The right panel shows the tracking results for the individual cells. The color and size of the arrow indicate migration velocity. The cyan and pink lines are outlines of the embryo at the first and second time points, respectively.

**Extended Data Fig. 4.**
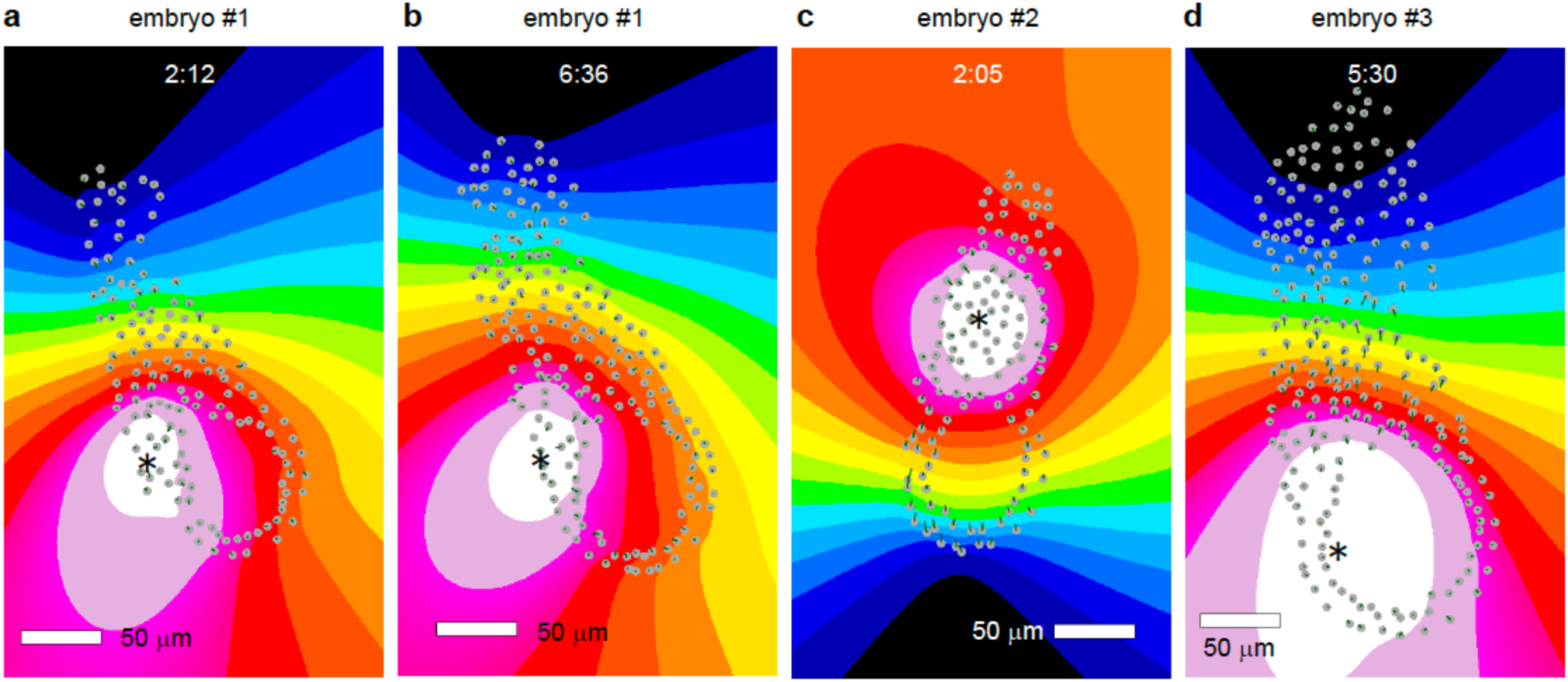
Shrinkage center of hiccup-like behavior in an E5.5 mouse embryo. (**a, b**) Visualization of shrinkage center of embryo #1 at time points 2:12 (**a**) and 6:36 (**b**) from the vertical view. (**c, d**) Visualization of shrinkage center of embryo #2 at time points 2:05 (**c**), and of embryo #3 at time points 5:30 (**d**) from the horizontal view. Colors indicates a shrinkage level at 4-bit scale, from white (min) to black (max). The gray circles are cell positions of the embryo and green lines indicates distances of cell migration within a frame. Asterisks show shrinkage center.

## Supplementary Notes

### Note 1. E5.5 mouse embryo observation using conventional microscopy

Before new construction, we performed test observations on the embryo of an *R26-H2B-EGFP* mouse line using four different types of commercialized microscopes: laser scanning confocal microscopy (LSCM), spinning-disk confocal microscopy (SDCM), two-photon excitation microscopy (TPEM), and multidirectional selective plane illumination microscopy (mSPIM) (Supplementary Fig. 1). Because our target was an elliptical sphere with a radius of 300 μm, the required observation depth was 300 μm. In addition, spatial resolution was required to individually isolate single cells. The observation depth did not reach 50 μm in the single-photon excitation of LSCM or SDCM. While TPEM could achieve an observation depth of more than 200 μm, its temporal resolution was insufficient. mSPIM only satisfied observation depth and temporal resolution, but not spatial resolution, to identify individual single cells owing to blurring by the lens effect and/or random light scattering by the microstructure in cells.

Photodamage in single-photon excitation confocal microscopy, including LSCM or SDCM, is a fatal problem for long-term embryo observation. The low temporal resolution of TPEM can be solved by the parallel use of spinning disk confocal microscopy.^29^ However, focusing a strong infrared laser is necessary to generate two-photon excitation, possibly causing a local temperature increase in the medium. The problem with the mSPIM application was the degradation in spatial resolution when observing deeper areas. To solve this problem, we constructed a dual-axes observation system, diSPIM, based on the mSPIM system.

We constructed an mSPIM system on a trial basis and investigated point spread functions (PSFs) along the x-, y-, and z-axes by observing fluorescent-labeled microspheres with a diameter of 0.2 μm (Fluoresbrite YG, Polysciences, Inc.), which were assumed as a point source, embedded in a 0.5% (w/v) agarose hydrogel. Even when using a sheet-formed laser, the resolution of the z-axis was significantly lower than those of the x- and y-axes (Supplementary Fig. 2a). The spatial resolution could be approximated using full width at half-maximum (FWHM) values obtained by fitting the fluorescent image of a microfluorophore with a Gaussian distribution (Supplementary Fig. 2b). The FWHMs of the x- and y-axes were 1.12 μm and 1.15 μm, respectively, and the FWHM of the z-axis was 5.08 μm at the just focused position. Furthermore, we noted that the resolution along the z-axis depended on the position along the beam propagation axis because of laser focusing (Supplementary Fig. 2c, *green*). Meanwhile, those of the x- and y-axes were independent of the horizontal position (Supplementary Fig. 2c, *red and blue*). Therefore, we used diSPIM.

### Note 2. Temperature stability via the two-layered incubator system

Our microscope system consisted of a two-layered incubator. A small chamber was installed on the microscope stage and the sample cuvette was mounted in the chamber (Fig. 1d, *orange*). The microscope, with the exception of the optics for laser illumination, was covered with a polycarbonate box (Supplementary Fig. 3, *right photograph*). The room temperature and relative humidity (RT) had to be maintained at 22.5 °C and < 40%, respectively, to maintain the performance of the equipment. The room air at 22.5 °C was heated to 37.5 °C and loaded into the box (Supplementary Fig. 3, *magenta*). A gas controller was used to adjust the CO_2_ concentration to 5% (Supplementary Fig. 3, *cyan*), and the 5% CO_2_ air, which obtained 95% RT via bubbling, was loaded into the chamber on the microscope (Supplementary Fig. 3, *orange*). In addition, the outer frame of the chamber had a groove for water to prevent evaporation of the medium (Supplementary Fig. 3, *upper right*).

The room temperature fluctuated within ±0.5 oC with a specific frequency (Supplementary Fig. 4a, *cyan*). The temperature of the medium in the cuvette synchronized the fluctuation up to 37.5 °C (Supplementary Fig. 4a, *magenta*). The polycarbonate box blocked the exchange of heat between the room and chamber, thus suppressing this temperature fluctuation (Supplementary Fig. 4b). Because of the small size of the chamber, the medium temperature in the cuvette was sensitive to heat transfer due to airflow caused by human operation (Supplementary Fig. 4c, *magenta*). The polycarbonate box also dramatically suppressed this (Supplementary Fig. 4c, *orange*). Moreover, evaporation of the medium was a critical problem for long-term observation; however, the two-layered incubator inhibited medium evaporation more than a commercialized incubator (Supplementary Fig. 4d).

### Note 3. Investigation of photodamage to embryonic stem cells (ESCs) during laser scanning

We developed a reproducible, quantitative, and easy evaluation method for photodamage utilizing the growth rate of the embryoid body (EB) before and after photo stimulus by focused-laser scanning as an index (Supplementary Fig. 5a, See method for details). An EB formed from 1000 cells over 48 h was damaged by laser scanning using a confocal microscope. Because the division rate of damaged cells decreases, the photodamage can be estimated from the ratio of the area of the EB before and 24 h after laser scanning. The actual measurement sometimes deviated from preconceived notions: after photo stimulus for 75 images, the growth rate correlated to stimulation photon energy excepting for the red laser, however, the red laser provided more damage than the green laser (Supplementary Fig. 5b, c). This result suggests that the cause of cell damage differs depending on wavelength.

Using the developed method, we screened the observation parameters involved in the photodamage before constructing the observation protocols. In this experiment, the laser wavelength was fixed at 488 nm because we used enhanced green fluorescent protein (eGFP), the most widely used fluorescent protein, as a model fluorescence tag. In general, photodamage is thought to depend on the laser irradiation power when using a single-color excitation. The growth rate of an EB depended on the laser intensity, as expected, however it was not linearly dependent; the growth rate decreased by reducing the laser intensity from 100% to 10%, decreased slightly by reducing this to 1%, and decreased again by reducing the intensity to 0.1% (Supplementary Fig. 6a). When setting laser intensity to 100% for the best image contrast, the parameters considered were scanning speed, frame number, and total exposure time in a photo stimulus by confocal scanning under constant laser output conditions. When fixing the total exposure time, the growth rate was negatively correlated with the scan speed (positively correlated with the frame number), despite the constant total photon energy received by the cells (Supplementary Fig. 6b). In short, when obtaining the same number of frames, the faster the scan speed, the lower the photodamage. Importantly, when the scanning speed was fixed, the reduction of photodamage by decreasing the frame rate saturated at the 5-min interval condition, and the growth rate did not change between the 5-min and 60-min intervals in this assay, even though the total number of photons changed by nine times (Supplementary Fig. 6c). This means that a smaller frame number does not necessarily reduce photodamage, and there is a best condition for minimum damage and maximum frame rate. Thus, we can determine the observation parameters to be searched for.

### Note 4. Investigation between photodamage and reactive oxygen species (ROS) production

Photodamage by laser irradiation has been attributed to reactive oxygen species (ROS) production.^15^ ROS production was estimated from the fluorescence intensity of an ROS-sensitive fluorescent probe (CellROX^R○^ DeepRed, Thermo Fisher C10422) introduced into cells (Supplementary Fig. 7a). ROS accumulated increasingly with each stimulus depending on the laser power (Supplementary Fig. 7b). Interestingly, the estimated ROS production rate was negatively correlated with the EB growth rate in laser power dependence (Supplementary Fig. 7c, *red, blue, green, and magenta*). Increasing the scan speed inhibited ROS production, even though the total number of frames acquired increased (Supplementary Fig. 7b, c, *orange*), which is consistent with the result that the faster the scan speed, the lower the photodamage (Supplementary Fig. 6b). The interval between two frames dramatically inhibited ROS production, whereas the growth rate did not recover by the interval (Supplementary Fig. 7c, *cyan, light green, and dark blue*), suggesting the existence of another cause of photodamage.

We expected a temperature increase inside an EB by strong laser irradiation, as previously reported,^30^ because the EB growth tested in this study was also affected by the temporal temperature increase in the culture medium (Supplementary Fig. 8). Intracellular temperature was estimated from the ratio of the fluorescent intensities of a dual-color thermoprobe (cationic fluorescent polymeric thermometers; Funakoshi FDV-0005) introduced into the cells (Supplementary Fig. 9a). The fluorescence ratio of the probe in an EB followed the medium temperature (Supplementary Fig. 9b). Subsequently, when estimating the temperature during the sequential acquisition of 55 frames with a scan speed of 4 s/frame, the temperature inside the EB changed slightly, even after entire acquisition (Supplementary Fig. 9c). It is unlikely that the temperature inside the EB increased to the extent that the growth was hindered by continuous imaging.

Thus, we surveyed the relationship between growth rate and ROS production and temperature increase. The ROS production correlated with the growth rate in a certain condition; however, it was not the only one. The mechanism of photodamage during microscopic observation includes not only ROS production but also many other mechanisms, such as direct attack on DNA or protein, the kinetics of ROS metabolism and recovery circuits, and damage to themselves. We will continue to investigate the mechanism of photodamage. Until the mechanism is clarified, there is no other way but to optimize it via actual measurement.

### Note 5. Construction of new optics for two-axes light sheets

Although the method using a cylindrical lens is the cheapest and simplest way to form the laser into a thin sheet, the sheet becomes barrel-shaped, resulting in an optical cross-section with spatially non-uniform thickness. To solve this problem, a method has been developed in which a focused laser beam is scanned in parallel to form a pseudo-light sheet with a uniform thickness along the scanning direction;^31^ many current SPIMs are based on this method. When the observation target is a biological sample, such as a mouse embryo, microstructures in cells, such as vesicles, partially cause scattering, refraction, and diffraction of the laser, resulting in stripe-shaped shadowing artifacts. mSPIM can reduce the artifacts by irradiating the light sheet from multiple directions.^7^ When collecting the image cross sections acquired using SPIM to reconstruct a 3D-image, the resolution difference between the in-plane (xy) and depth (z) directions is a problem (Supplementary Fig. 2). The image cross-sections (xz or yz) on the second axis can be acquired by rotating the sample by 90°, which is time consuming. Subsequently, diSPIM was developed to switch between an xy-plane and a plane perpendicular to it (xz or yz) using a two-dimensional scanner.^8^

For the parallel use of mSIPM and diSPIM, two 4f imaging systems, in which Galvano mirrors (GMs) are set on conjugate focal planes and conjugate pupil planes, should be arranged in series on one optical path (Supplementary Fig.10a). However, it is practically difficult to produce the exact same light sheets on both the horizontal and vertical axes; one light sheet is generated by a GM in the former 4f system (Supplementary Fig. 10a, *GM2*), whereas the other light sheet is generated by a GM in the latter 4f system (Supplementary Fig. 10a, *GM4*). The influence of various aberrations of the relay lens on the optical path cannot be ignored. The order of the GM for controlling the sheet height (focal plane) and the GM for sheet forming is reversed between the orthogonal light sheets; therefore, the orthogonal light sheets are not optically equivalent. Hence, we developed a new optical system in which the optics of the horizontal and vertical sheets are separated and arranged in parallel (Supplementary Fig. 10b).

Our application was required to adjust the waist position of the light sheet to the position of each sample. For this purpose, we installed an electrically focus-tunable (varifocal) lens on the conjugate pupil plane (Supplementary Fig. 10b, *VL*), which changed the divergence of the laser beam, to adjust the focus position of the laser. Galvanometric mirrors (Supplementary Fig. 10b, *GM1 and GM2*) were placed on the conjugate focal plane and the following pupil plane after VL. VL and GM2 were relayed using a 4f system consisting of a pair of convex lenses (Supplementary Fig. 10b, *L1* and *L2*). The other 4f system consisting of another pair of convex lenses (Supplementary Fig. 10b, *L3* and *L4*) was constructed such that a galvanometric mirror (Supplementary Fig. 10b, *GM3*) was located on the pupil plane. Subsequently, we constructed a bifurcated 4f system consisting of three convex lenses (Supplementary Fig. 10b, *L5*, *L6*, and *L7* and a polarized beam splitter (Supplementary Fig. 10b, *PBS1*). The bifurcated 4f system generates orthogonal light sheets in the optical paths (Supplementary Fig. 10b, *P1 and P2*). The generated light sheets were merged onto the same pathway using a half-wave plate (Supplementary Fig. 10b, *HWP*) and another polarized beam splitter (Supplementary Fig. 10b, *PBS2*). Each of the 4f systems consisted of the lens pairs (L5, L7) and (L6, L7), which relayed the pupil plane at GM3 to the backfocal plane of an objective lens to form a light sheet into the sample. Optical beam shutters were installed on each of the branched paths (Supplementary Fig. 10b, *S1 and S2*). A light sheet was generated by scanning GM2. Scanning GM1 resulted in the translation of the laser beam incident on GM2, realizing mSPIM. GM3 was used to control the position (height) of the light sheet perpendicular to it. This allowed the two orthogonal light sheets bifurcated by PBS1 to be optically equal and alternately switched by S1 and S2. This optical system requires a coherent and nonpolarized or circularly polarized laser. All optical parts including lenses, slits, PBSs, and HWP were purchased by Thorlabs. VL and GMs were purchased by Optotune and Cambridge Technology respectively.

### Note 6. Limitation of the present system for in-toto single-cell observation in an E5.5 mouse embryo

Photodamage determines the success or failure of long-term single-cell observation in a living E5.5 mouse embryo. To confirm that the embryos we observed were developing normally, we must determine the phenotype of a mouse embryo with photodamage. Because the higher power of laser irradiation caused more severe photodamage to EBs as a lower growth ratio (Supplementary Fig. 6a), we performed time-lapse imaging of an E5.5 embryo irradiated with ten times higher irradiation power than under normal conditions. Dead cells were observed in the epiblast within 1 h of time-lapse imaging (Supplementary Fig. 11, *left panels*, Supplementary Videos 7 and 8), cell debris gradually accumulated in the amniotic cavity, and the embryo stopped growing and became small (Supplementary Fig. 11, *arrows*). The extraembryonic ectoderm appeared to be damaged. The increase in the overall embryo volume was significantly reduced 3 h after the start of observation. However, in the visceral endoderm, cell division was observed, and cell death was not remarkable, even 5 h after the start. These results suggest that the sensitivity to photodamage differs among various tissues, and the epiblast is most sensitive at the E5.5 stage. It is also possible that the differentiating state affects sensitivity to photodamage.

The two-layered incubation and laser irradiation protocol enabled the continuous observation of a living mouse E5.5 embryo every 5–6 min for more than 15 h and sometimes more than 24 h. However, although diSPIM can compensate for the elongation of the PSF along the optical axis on one orthogonal plane using the other plane, our optical system could not improve image degradation in deep biological areas because we used only linear optics. Up to 5–6 h after the E5.5 stage, all the cells could be reliably isolated using 3D stacks of the images of either plane or 3D rendered images constructed from the dual-axes image stacks (Supplementary Fig. 12, *left and second left*). After 10–12 hours, it was difficult to isolate single cells with only a snapshot (Supplementary Fig. 12, *second right*). However, single-cell isolation was possible by the human eye with reference to temporal information. Single-cell isolation was still possible in the ectoplacental cone and extraembryonic visceral endoderm, but some cells in the extraembryonic ectoderm could no longer be isolated (Supplementary Fig. 12, *right*). In short, we conclude that the present system enables in-toto single-cell tracking in a hemispherical E5.5 embryo, limited to 12 h. When entire embryo observation is required, the other hemispherical side can be observed by installing an additional camera. Notably, we achieved single-cell resolution in the entire E5.5 embryo despite the adverse optical conditions of blue laser excitation. Using green-excited red fluorescent proteins, such as mCherry and mKate,^32, 33^ instead of GFP, is expected to reduce phototoxicity and improve the low permeability of light.

---

**Supplementary Fig. 1.**
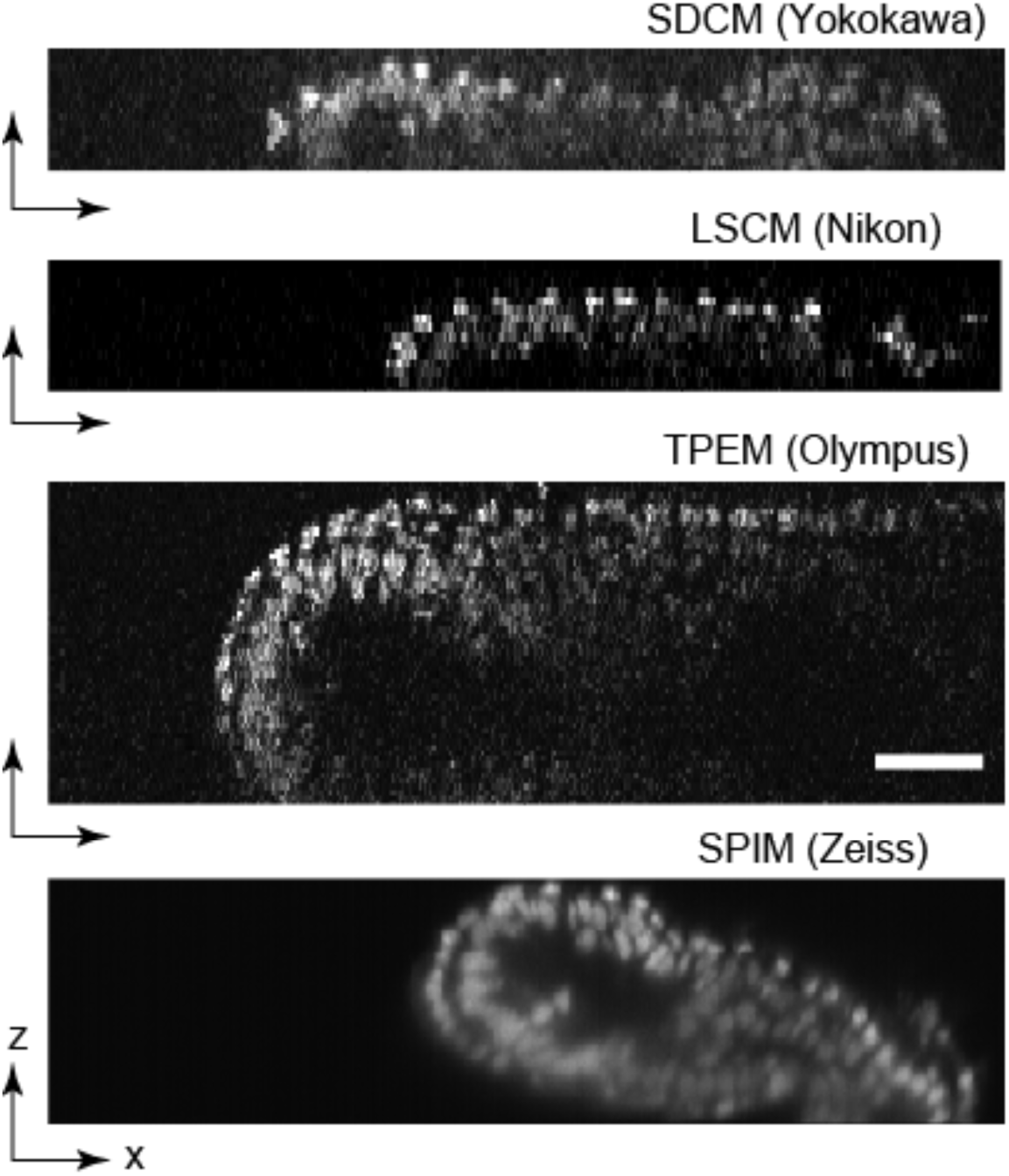
Comparison of E5.5 or E6.5 mouse embryo observations in four branches of conventional microscopy. Images of mouse embryos expressing H2B fused with green fluorescent protein (H2B-GFP) were obtained by spinning-disk confocal microscopy (SDCM; CV1000, Yokogawa), laser scanning confocal microscopy (LSCM; A1, Nikon), two-photon excitation microscopy (TPEM; FV1000-MPE, Olympus), and multidirectional selective plane illumination microscopy (mSPIM; Z1, Zeiss). In the case of TPEM, an E6.5 embryo was imaged because the observation depth was easily achieved to observe the entire E5.5 embryo. In other cases, an E5.5 embryo was used. The scale bar is 100 μm.

**Supplementary Fig. 2.**
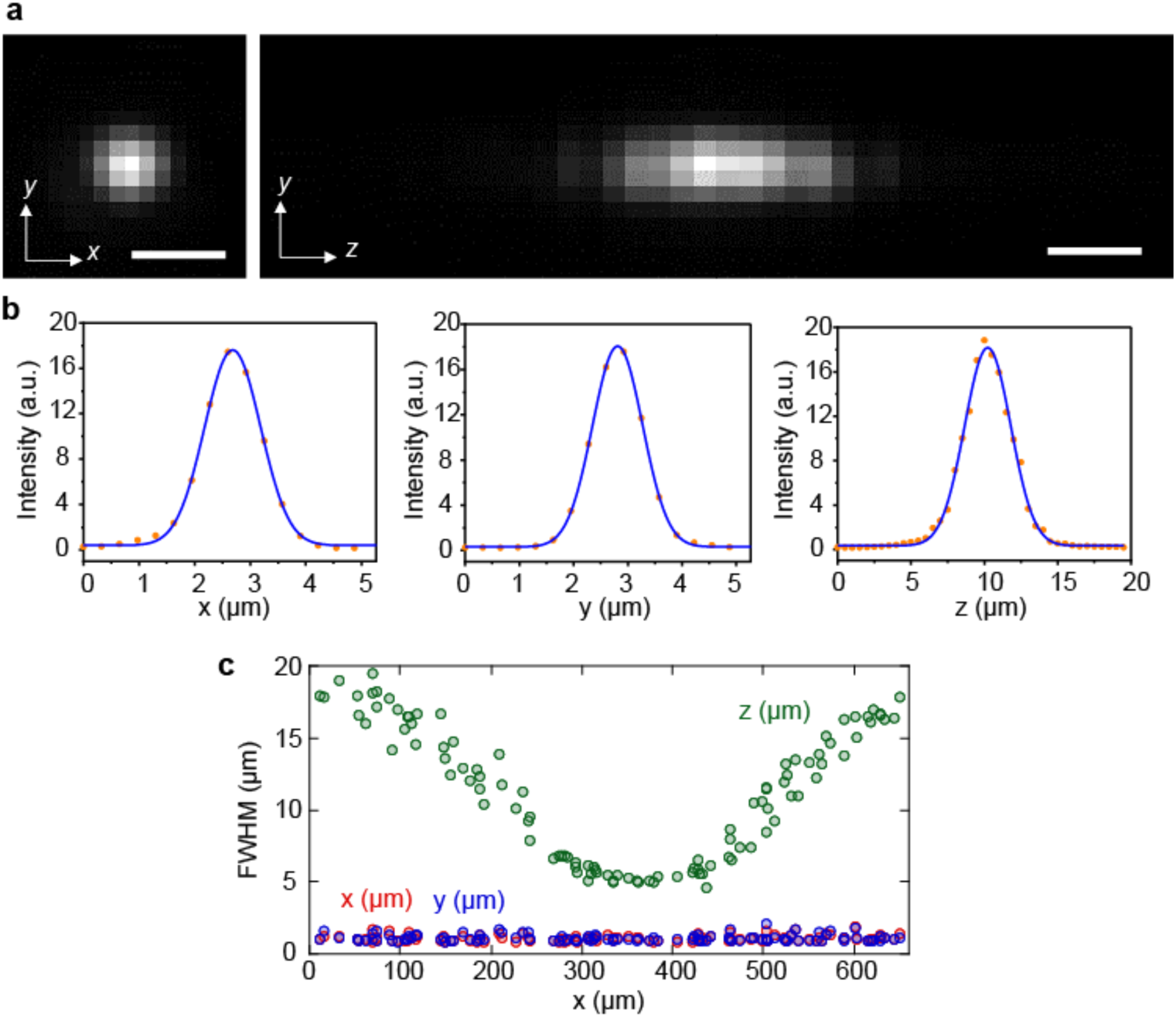
Point spread function measured using mSPIM. (**a**) Fluorescent images of fluorescently labeled microspheres with a diameter of 0.2 μm (Fluoresbrite YG, Polysciences, Inc.) on the y–x plane (*left*) and y–z plane (*right*) obtained using homemade mSPIM. The microspheres were embedded in a 0.5% (w/v) agarose hydrogel. Images were taken at a depth of 0.5–0.6 mm from the glass surface. The z-position was stepped every 0.5 μm for 200 image acquisitions (corresponding axial (z) range of 100 μm). The number density of the microspheres in the sample was 7.4 × 108 mL^-1^. (**b**) Intensity profiles across a fluorescent microsphere along the x-(*left*), y-(*center*), and z-axes (*right*). The solid lines are the fitting results with the Gaussian functions, which are used to obtain the full width at half-maximum (FWHM) values of the fluorescence peak around the microsphere. (**c**) Position dependency of the FWHM of the x-(*red*), y-(*blue*), and z-axes (*green*).

**Supplementary Fig. 3.**
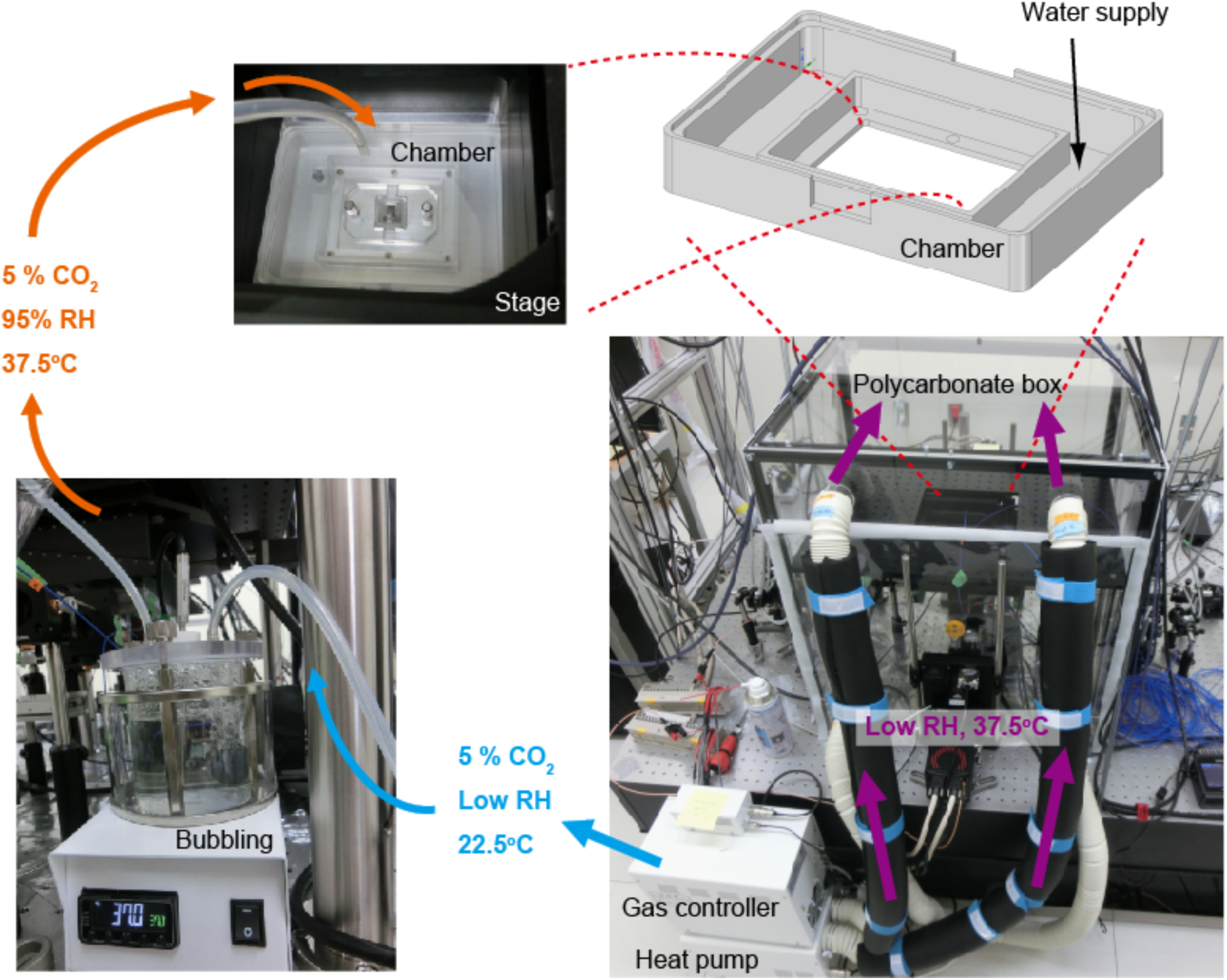
Two-layered incubation system. Schematic illustration and photographs of the system of air heating, bubbling, and loading. Air with 5% CO_2_, 95% relative humidity (RH), and a temperature of 37.5 °C was loaded into the home-made chamber on the microscope via a gas controller and bubbling device. The outer frame of the chamber contained grooves for water, and air of 37.5 °C was loaded into the polycarbonate box covering the microscope.

**Supplementary Fig. 4.**
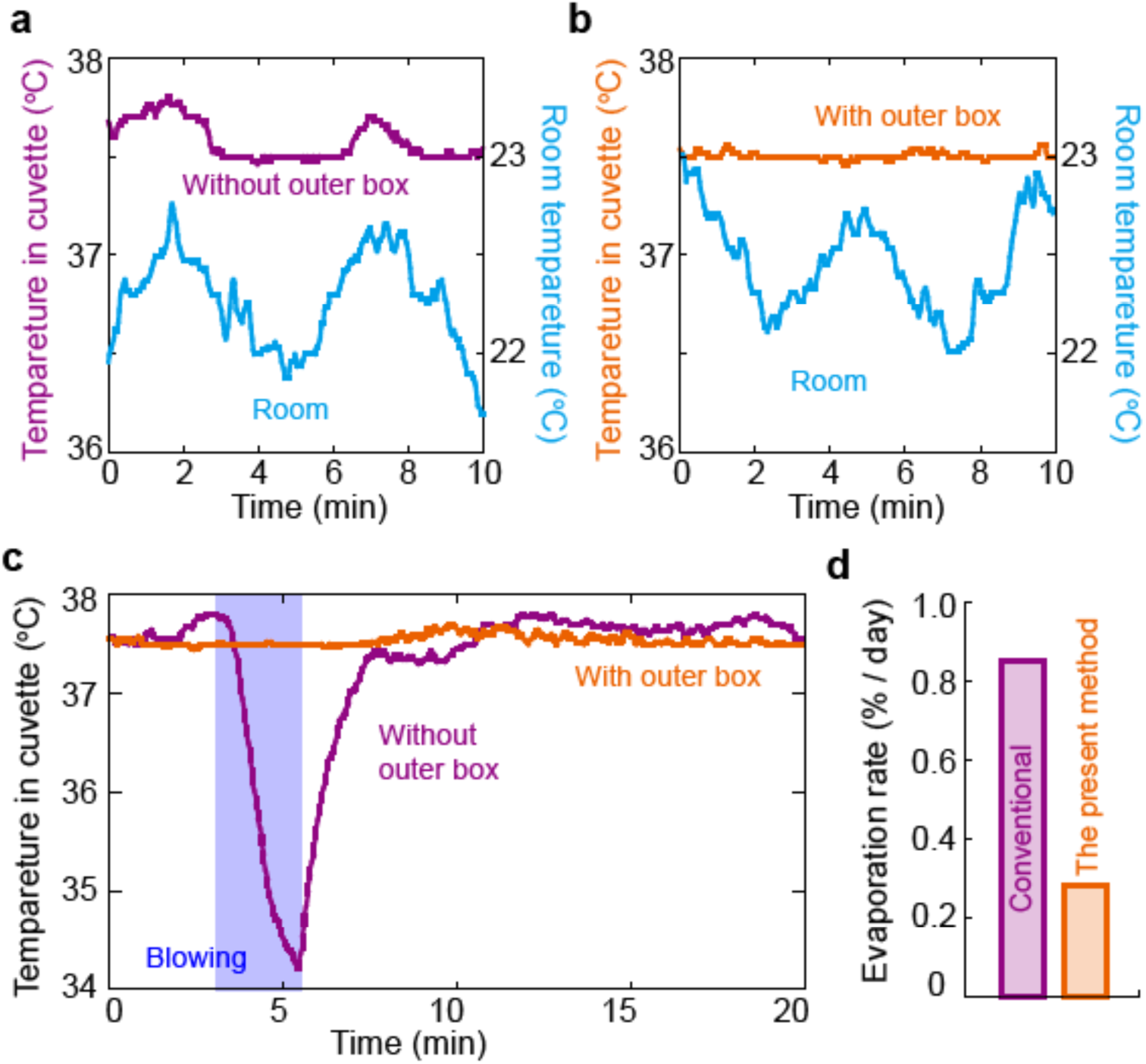
Temperature stabilization using the two-layered incubation system. (**a, b**) Time course of the temperature of the medium inside the cuvette without (**a**) and with (**b**) the polycarbonate box. Cyan, room temperature; magenta or orange, medium temperature. (**c**) Time course of the temperature of the medium inside the cuvette without (*magenta*) and with (*orange*) the polycarbonate box when blowing due to human behavior (*blue*). (**d**) Medium evaporation rate of 200 μL was measured in a commercial conventional incubator (*magenta*, Astec APM-30DR) and in the two-layered incubation system (*orange*).

**Supplementary Fig. 5.**
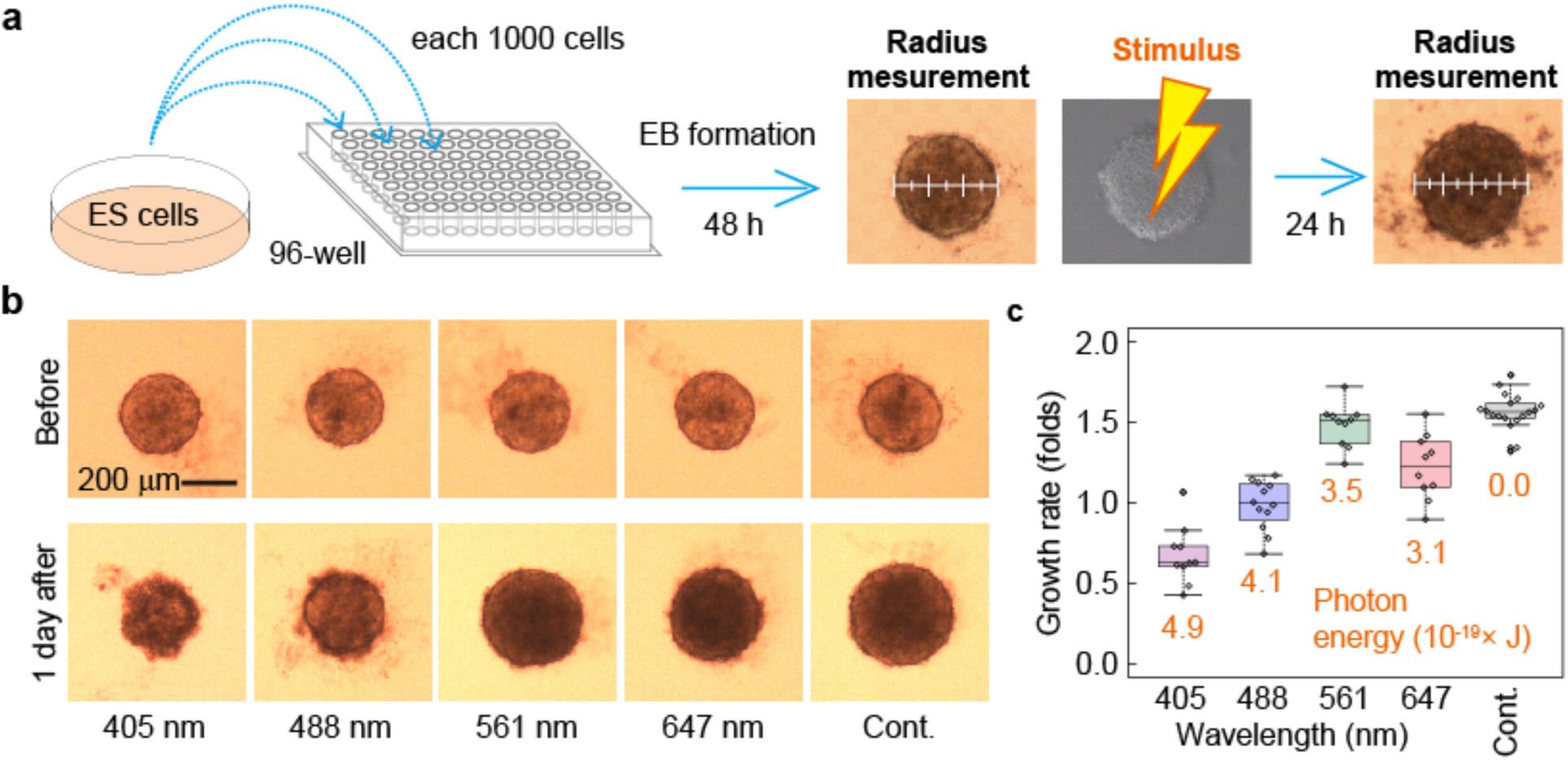
Assay system to investigate photodamage using embryoid body (EB) growth. (**a**) Explanation of the assay protocol to investigate EB photodamage. The growth rate of an EB can be estimated from the ratio of the area of the EB before and after photo stimulus using confocal scanning. (**b)** Example of photodamage assay and excitation dependence. Bright-field images of each EB were recorded before (*upper*) and 1 d after (*lower*) photo stimulus via confocal scanning (75 times with a scan speed of 4 s/frame without intervals). (**c**) Box plot of the EB growth rate defined by the ratio of EB diameters before and 1 d after stimulation with various wavelengths. The values in the graph indicate the photon energy received by an EB due to photo stimulus. Each circle indicates a single EB.

**Supplementary Fig. 6.**
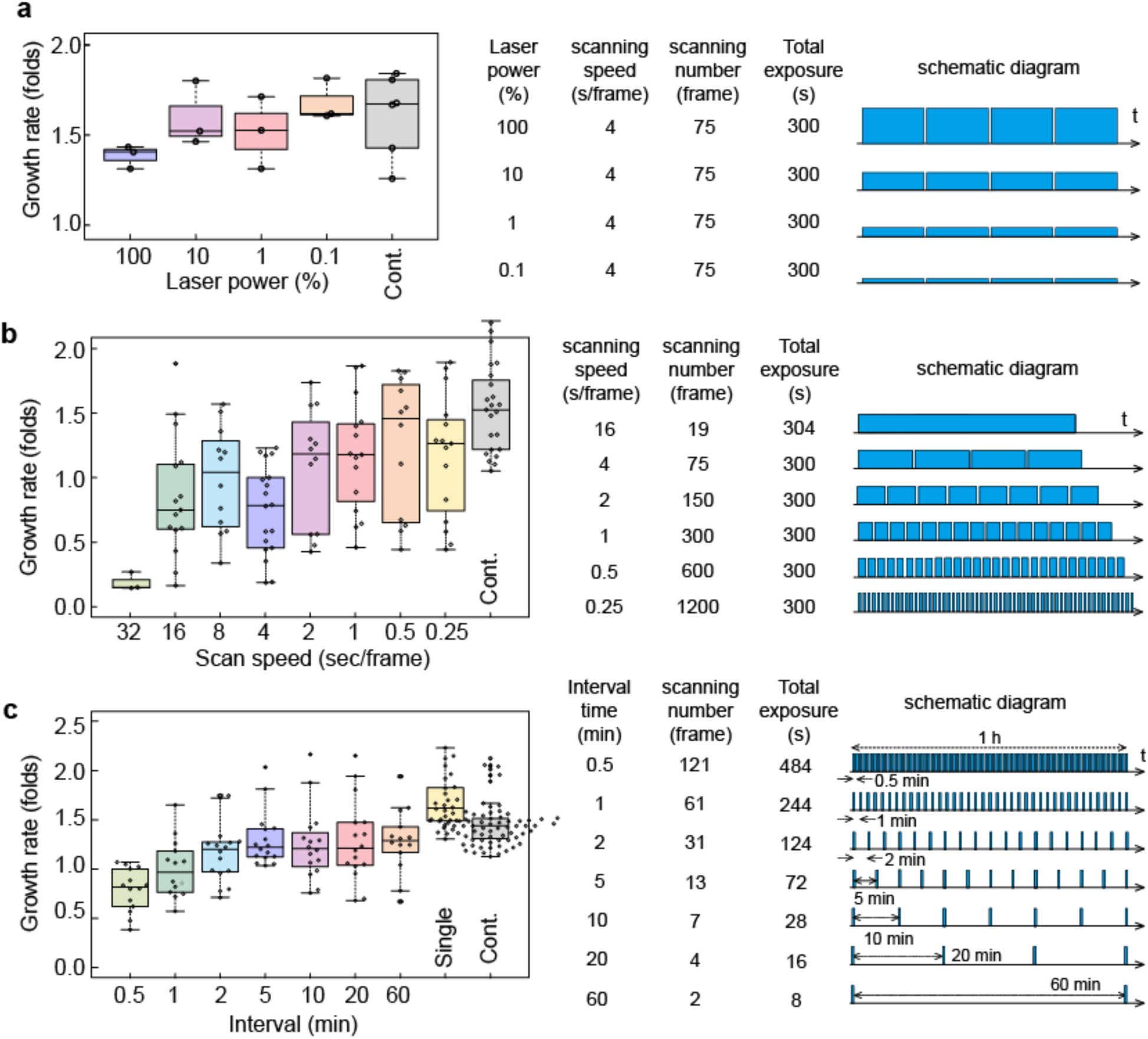
Investigation of photodamage on EB growth. Box plots of scan speed dependence on excitation power dependence (**a**), growth rate (**b**), and interval time between two frames (**c**). Right, estimated growth rates; left, conditions and schematic diagram of each assay. For **b**, the scan speed was fixed at 4 s/frame. Each circle indicates a single EB. The control (*Cont.*) was data stimulated by the same scanning procedure with 0% output of the laser (*gray*).

**Supplementary Fig. 7.**
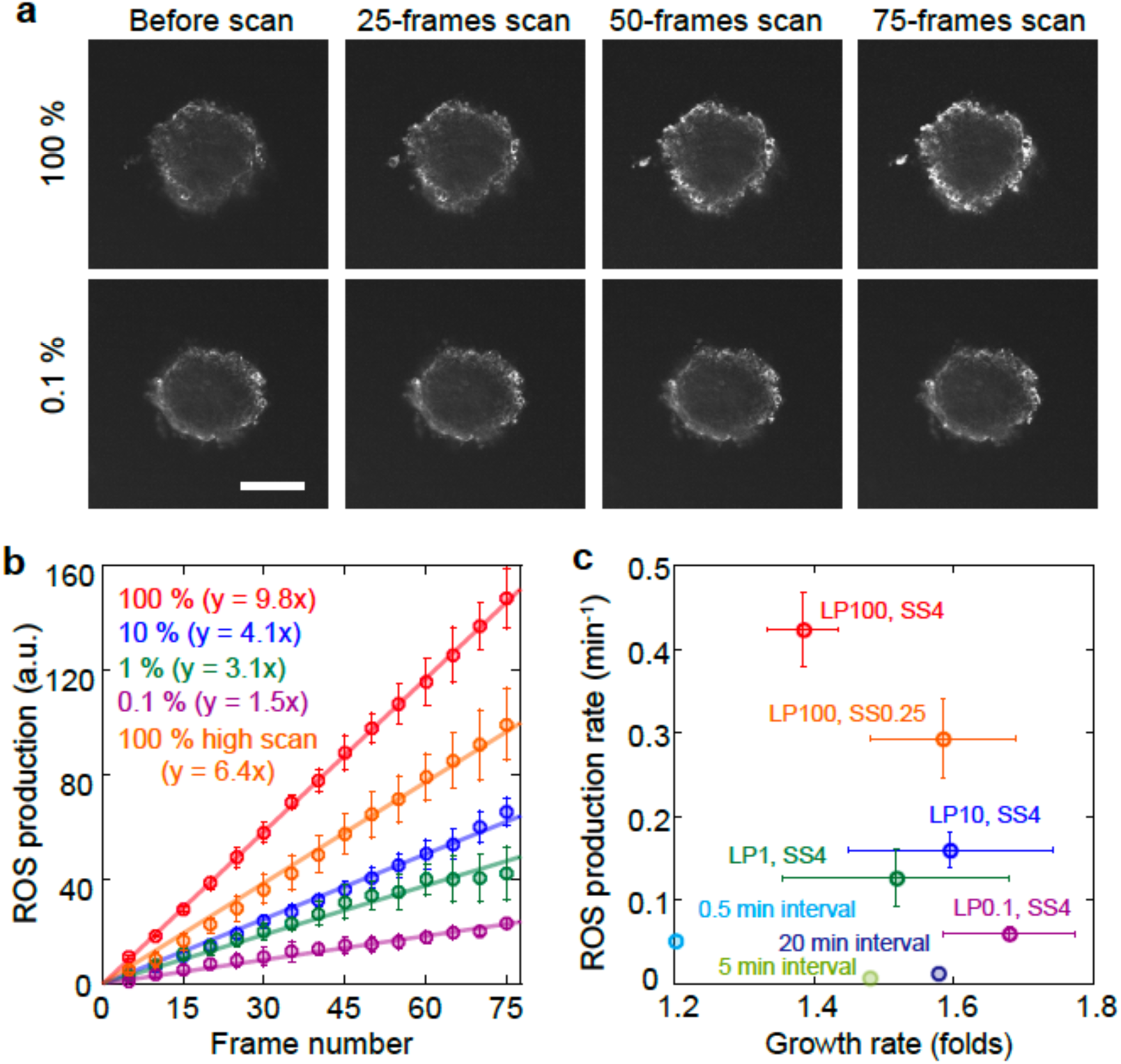
Evaluation of reactive oxygen species (ROS) production in an EB during confocal fluorescent imaging. (**a**) Time course of confocal fluorescent images of ROS sensitive dye in an EB before and after image excitation by 100% output (*upper*) and 0.1% output (*lower*) of a 488 nm laser. The image contrast was normalized to that before scanning (*left*). The scale bar is 100 μm. (**b**) Time course of estimated ROS production during confocal fluorescence imaging with a scan speed of 4 s/frame excited by 100% (*red*), 10% (*blue*), 1% (*green*), and 0.1% (*magenta*). The slope of the linear approximation indicates the ROS production rate (*lines*). (**c**) Correlation between the ROS production rate and the EB growth rate during confocal fluorescence imaging with a scan speed of 4 s/frame excited by 100% (*red*), 10% (*blue*), 1% (*green*), and 0.1% (*magenta*). The data from varying the interval period, that is, 0.5 (*cyan*), 5 (*light green*), and 20 min (*dark blue*), when setting the scan speed to 4 s/frame and excitation laser power to 100% overlap in the same graph; however, this was the result of only one trial. The error bars represent the standard deviation. LP, excitation laser power; SS, scan speed. The data when setting the scan speed to 0.25 s/frame and excitation laser power to 100% overlap in **b** and **c** (*orange*).

**Supplementary Fig. 8.**
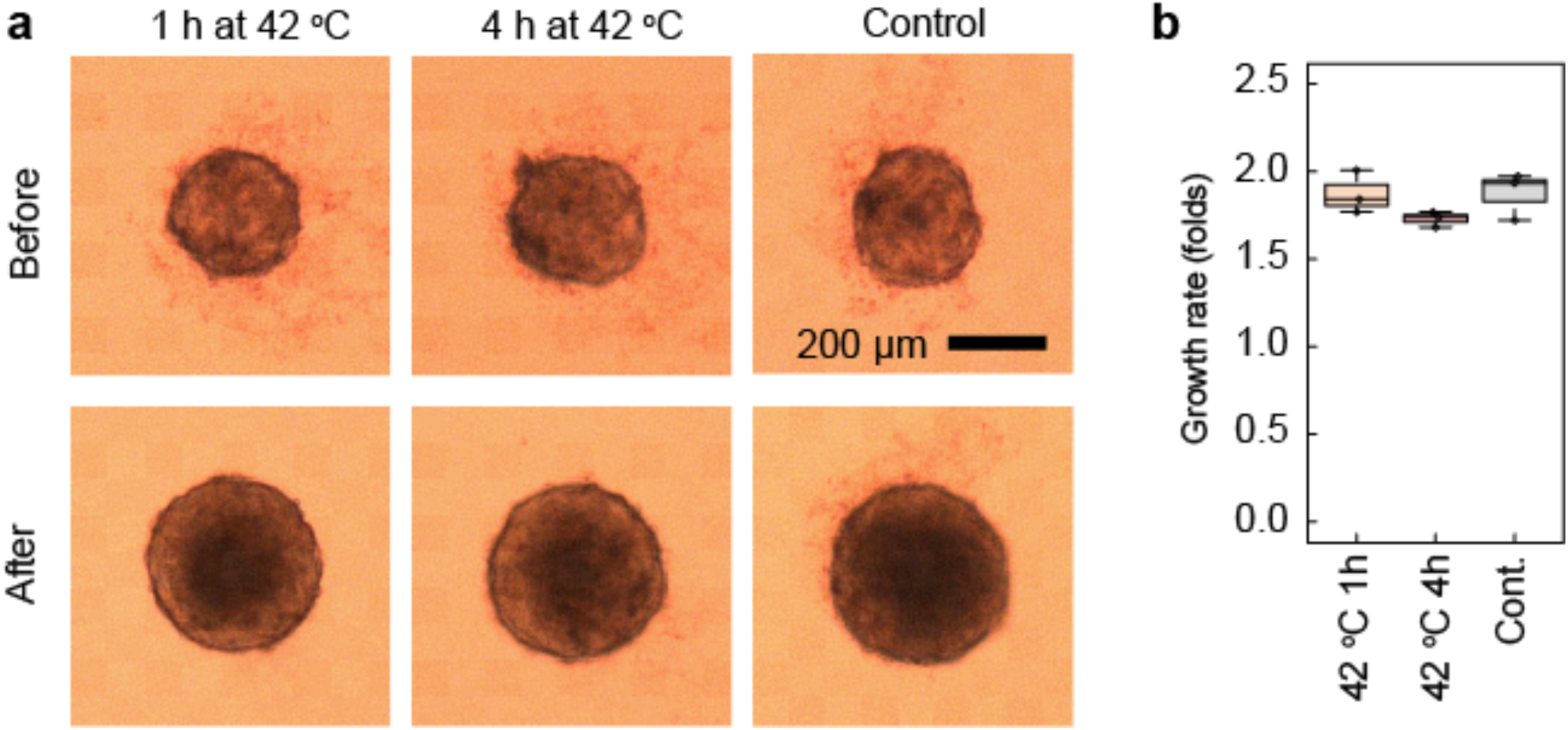
Temperature dependence of EB growth rate. (**a**) Bright-field images of an EB before (*upper*) and after (*lower*) temperature stimulus instead of photo stimulus. After EB formation, the EB was cultured for 24 h in conditions of 37 °C, 5% CO_2_, and >95% RH (*right*). The temperature was increased to 42 °C for the first 1 h (*left*) or 4 h (*middle*) of the 24 h period. (**b**) Box plots of the estimated EB growth rate after the 1 h temperature increase (*left*), 4 h temperature increase (*middle*), and no temperature increase (*right*).

**Supplementary Fig. 9.**
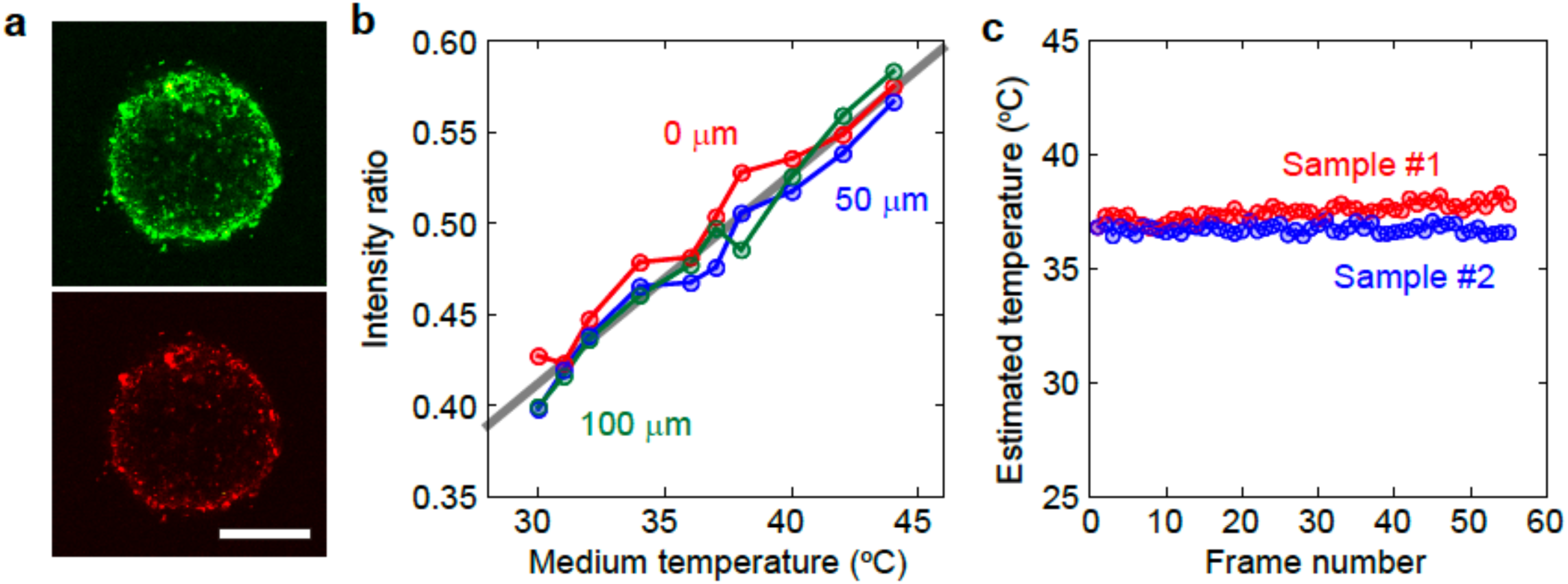
Measurement of intracellular temperature in an EB during fluorescent imaging. (**a**) Fluorescent images of an EB labeled with a dual-color thermoprobe (cationic fluorescent polymeric thermometers). The thermoprobe included two fluorophores: DBThD-AA and BODIPY-AA. The emission intensity of DBThD-AA (*upper*), emitting green fluorescence, decreases with increasing temperature in solution, while that of BODIPY-AA, emitting red fluorescence (*lower*), does not depend on the solution temperature. The scale bar is 200 μm. (**b**) Dependence of the intensity ratio of DBThD-AA fluorescence and BODIPY-AA fluorescence on solution temperature at the surface of an EB (*red*), a depth of 50 μm (*blue*), and a depth of 100 μm (*green*) in the EB. The temperature inside the EB was estimated using the calibration curve obtained from the plots (*gray line*). (**c**) Typical time course of the temperature at a depth of 100 μm in an EB during sequential image acquisition with a scan speed of 4 s/frame and excitation laser power of 100%.

**Supplementary Fig. 10.**
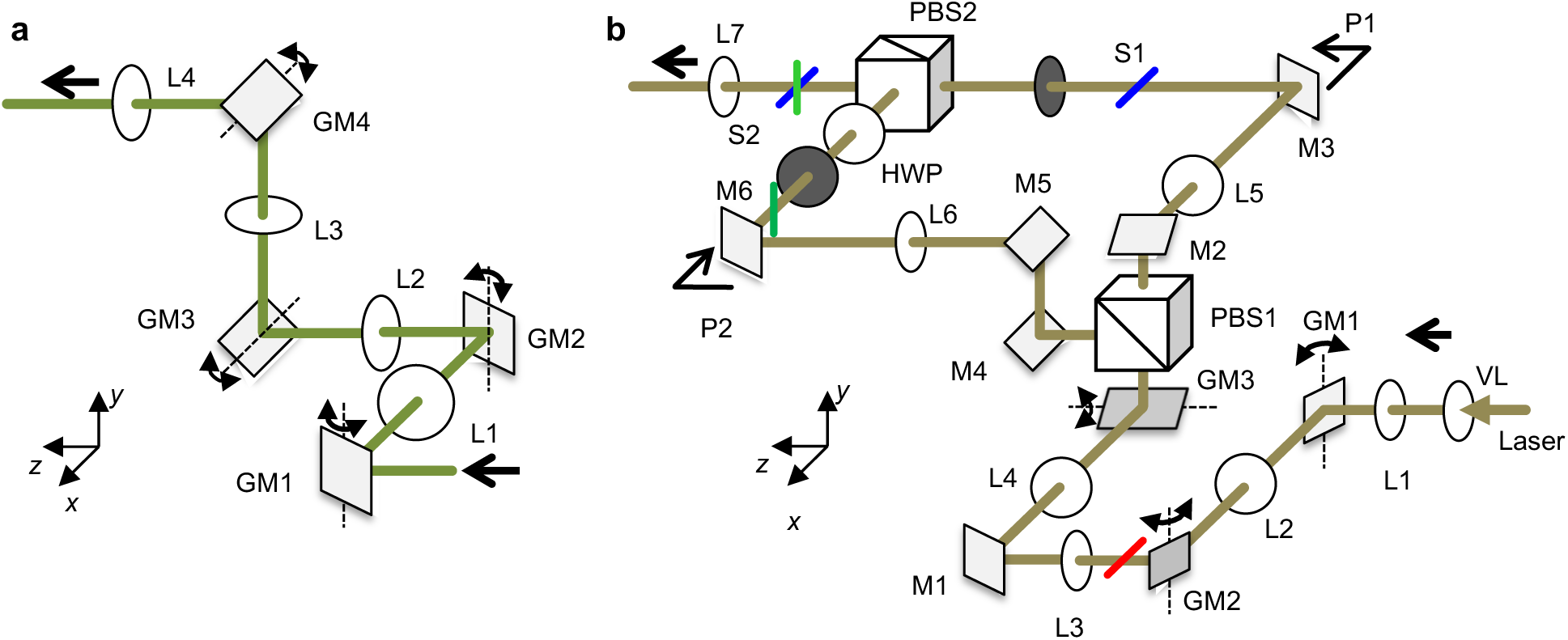
Optical setup for parallel use of mSPIM and diSPIM. (**a**) Implementation of an optical system with successive optical elements to generate two orthogonal light sheets. GM, galvanometric mirror; L, plano-convex lens. GM1 and GM3 are located on conjugate focal planes, whereas GM2 and GM4 are located on conjugate pupil planes. (**b**) Optical system with a bifurcated 4f system developed and used in the present study instead of (**a**), symmetrically placed on both sides of the incubation box in Fig. 1(b). VL, varifocal lens; L, plano-convex lens; GM, galvanometric mirror; M, dielectric multilayer mirror; PBS, polarized beam splitter; HWP, half-wave plate; S, mechanical shutter. GM1 is located on a conjugate focal plane, whereas VL, GM2, and GM3 are located on pupil planes. The red, blue, and green solid lines represent the scanning directions of the laser beam at each point.

**Supplementary Fig. 11.**
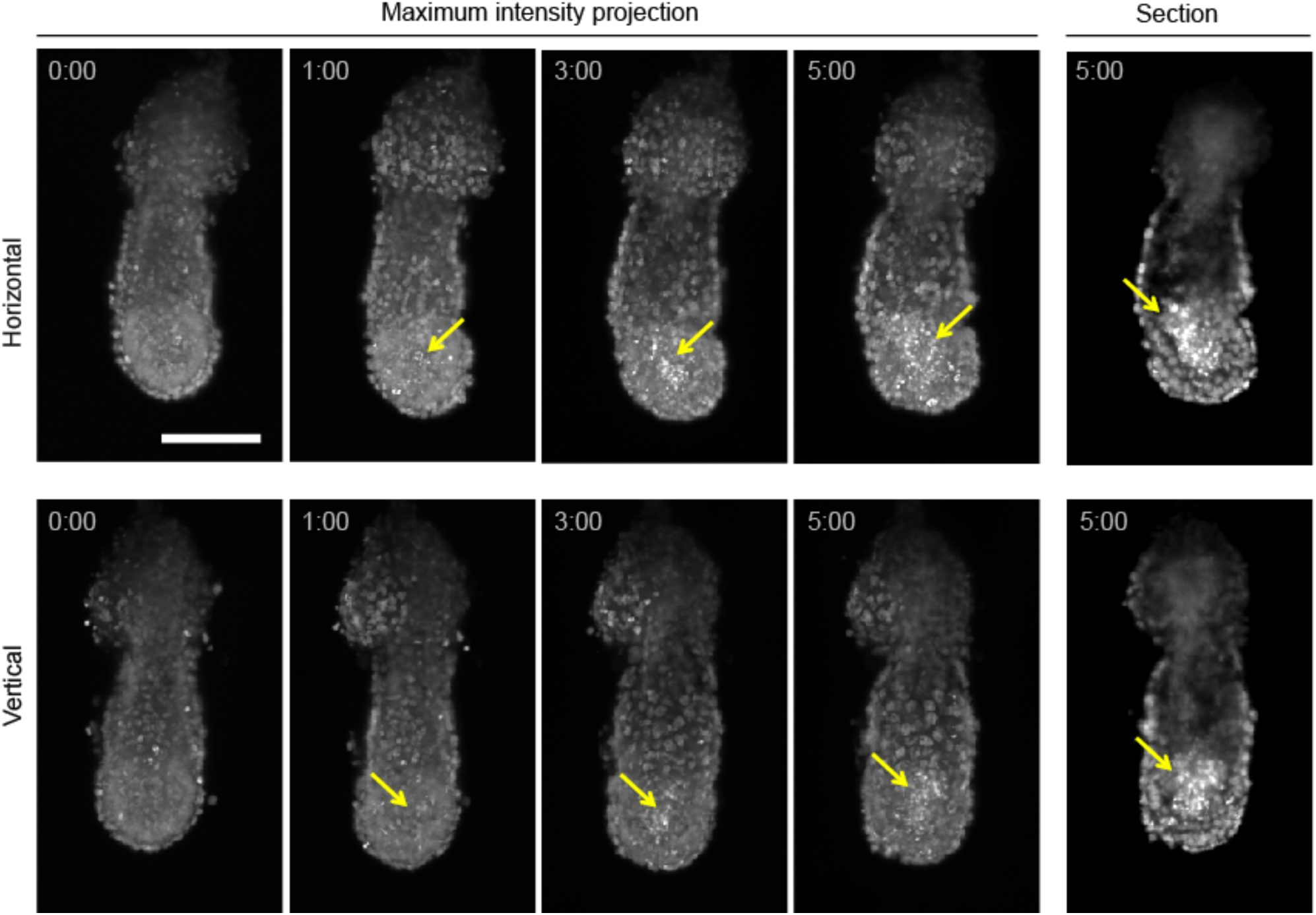
Time-lapse images of a mouse E5.5 embryo damaged by high-power laser irradiation. Maximum intensity projection of the time-lapse images of an *R26-H2B-EGFP* mouse embryo at E5.5. Sixty-one horizontal and 71 vertical images with a 5 μm z-step were acquired with high-power laser excitation every 5 min for 5 h. Cell debris (yellow arrows) appeared in the proamniotic cavity within 1 h of the start time and increased during time-lapse imaging. The scale bar is 100 μm.

**Supplementary Fig. 12.**
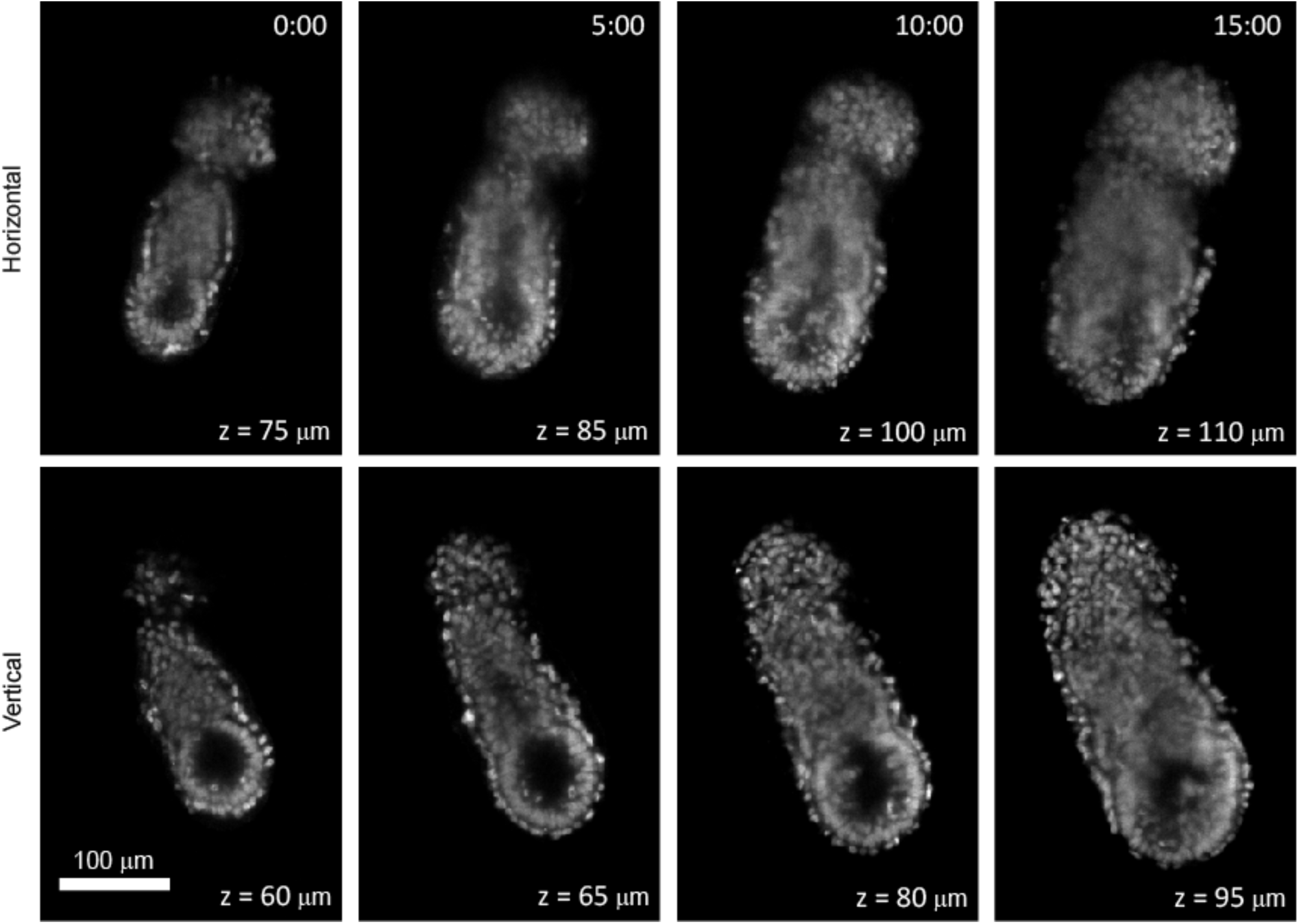
Time-lapse images of a mouse E5.5 embryo in the deepest region. Optical sections of the time-lapse images of an *R26-H2B-EGFP* E5.5 mouse embryo in the deepest region. The scale bar is 100 μm. The z-value represents the observation depth.

## Supplementary Videos

### Supplementary Video 1

Maximum intensity projection of the time-lapse images of an *R26-H2B-EGFP* mouse embryo at E5.5, related to Fig. 1g. Images were acquired every 6 min for 15 h.

### Supplementary Video 2

Optical sectional time-lapse images of the *R26-H2B-EGFP* mouse embryo at E5.5, related to Supplementary Video 1. Images were acquired every 6 min for 15 h.

### Supplementary Video 3

Maximum intensity projection of the time-lapse images of *R26-H2B-EGFP* mouse embryo #2 at E5.5, related to Fig. 2e. Images were acquired every 5 min for 15 h.

### Supplementary Video 4

Optical sectional time-lapse images of *R26-H2B-EGFP* mouse embryo #2 at E5.5, related to Supplementary Video 3. Images were acquired every 6 min for 15 h.

### Supplementary Video 5

Maximum intensity projection of the time-lapse images of *R26-H2B-EGFP* mouse embryo #3 at E5.5, related to Fig. 2e. Images were acquired every 6 min for 15 h.

### Supplementary Video 6

Optical sectional time-lapse images of *R26-H2B-EGFP* mouse embryo #3 at E5.5, related to Supplementary Video 5. Images were acquired every 6 min for 15 h.

### Supplementary Video 7

Maximum intensity projection of the time-lapse images of the *R26-H2B-EGFP* mouse embryo at E5.5, damaged by high-power laser irradiation. Images were acquired every 5 min for 20 h.

### Supplementary Video 8

Optical sectional time-lapse images of the *R26-H2B-EGFP* mouse embryo at E5.5, damaged by high-power laser irradiation. Images were acquired every 5 min for 20 h.

### Supplementary Video 9

Cell tracking images of the *R26-H2B-EGFP* mouse embryo at E5.5, related to Fig. 2d, from the lateral view. Individual cells were tracked from the elapsed time 0:00 to 6:00. A 300 min subset of cell tracks is shown by color lines. Green, DVE; blue, visceral endoderm; yellow, epiblast.

### Supplementary Video 10

Cell tracking images of the *R26-H2B-EGFP* mouse embryo at E5.5, related to Fig. 2d, from the anterior view. Individual cells were tracked from the elapsed time 0:00 to 6:00. A 300 min subset of cell tracks is shown by color lines. Green, DVE; blue, visceral endoderm; yellow, epiblast.

### Supplementary Video 11

Cell tracking images of the *R26-H2B-EGFP* mouse embryo at E5.5, related to Fig. 2d, from the distal view. Individual cells were tracked from 0:00 to 6:00. A 300 min subset of cell tracks is shown by color lines. Green, DVE; blue, visceral endoderm; yellow, epiblast.

### Supplementary Video 12

Maximum intensity projection of the time-lapse images of the *R26-H2B-EGFP* mouse embryo at E6.5, related to Fig. 1f. Images were acquired every 5 min for 18 h.

### Supplementary Video 13

Optical sectional time-lapse images of the *R26-H2B-EGFP* mouse embryo at E6.5, related to Supplementary Video 12. Images were acquired every 5 min for 18 h.

